# Hybrid computational modeling highlights reverse Warburg effect in breast cancer-associated fibroblasts

**DOI:** 10.1101/2023.05.11.540378

**Authors:** Sahar Aghakhani, Sacha E Silva-Saffar, Sylvain Soliman, Anna Niarakis

## Abstract

Cancer-associated fibroblasts (CAFs) are key players of the tumor microenvironment (TME) involved in cancer initiation, progression, and resistance to therapy. These cells exhibit aggressive phenotypes affecting, among others, extracellular matrix remodeling, angiogenesis, immune system modulation, tumor growth, and proliferation. CAFs phenotypic changes appear to be associated with metabolic alterations, notably a reverse Warburg effect that may drive fibroblasts transformation. However, its precise molecular mechanisms and regulatory drivers are still under investigation. Deciphering the reverse Warburg effect in breast CAFs may contribute to a better understanding of the interplay between TME and tumor cells, leading to new treatment strategies. In this regard, dynamic modeling approaches able to span multiple biological layers are essential to capture the emergent properties of various biological entities when complex and intertwined pathways are involved. This work presents the first hybrid large-scale computational model for breast CAFs covering major cellular signaling, gene regulation, and metabolic processes. It was generated by combining an asynchronous cell- and disease-specific regulatory Boolean model with a generic core metabolic network leveraging both data-driven and manual curation approaches. This model reproduces the experimentally observed reverse Warburg effect in breast CAFs and further identifies Hypoxia-Inducible Factor 1 (HIF-1) as its key molecular driver. Targeting HIF-1 as part of a TME-centered therapeutic strategy may prove beneficial in the treatment of breast cancer by addressing the reverse Warburg effect. Such findings in CAFs, considering our previously published results in rheumatoid arthritis synovial fibroblasts, point to a common HIF-1-driven metabolic reprogramming of fibroblasts in breast cancer and rheumatoid arthritis.

All analyses are compiled and thoroughly annotated in Jupyter notebooks and R scripts available on a GitLab repository (https://gitlab.com/genhotel/breast-cafs-reverse-warburg-effect) and a Zenodo permanent archive [1].

## 1 Introduction

Cancer is the second leading cause of mortality worldwide, accounting for nearly 10 million deaths in 2020 [2]. Recognition of its burden on patients, families, communities, and health care systems as a major public health concern boosted research on the disease in an unprecedented way. Over the past two decades, numerous prevention strategies have been developed to avoid major risk factors, including screening and early detection protocols, successfully preventing an estimated 30-50% of cancer-related deaths [2]. Additionally, a variety of treatment options are currently available including surgical interventions, chemotherapies, radiotherapies, immunotherapies, hormone therapies, targeted drug treatments, etc. However, a significant part of drug-based therapeutic approaches still fails, mainly due to drug resistance, representing the main challenge of cancer researchers to date. Resistance can occur when cancer cells present molecular alterations driving insensitivity to drugs before treatment, called intrinsic resistance, or when cancer cells adapt to a drug while it is being administered, known as extrinsic resistance. The latter is suspected to arise from changes in the tumor microenvironment (TME), as its interplay with cancer cells is often partially or totally overlooked in the development of cancer therapy.

Yet, the importance of the TME in cancer initiation and progression has been widely recognized for years [3-5]. While cancer develops through genetic and epigenetic alterations, tumor growth, survival, and metastasis are regulated through complex interactions with stromal cells of the TME [6]. Among them, multiple studies in various cancers have demonstrated the key role of cancer-associated fibroblasts (CAFs) [7-9]. Quiescent fibroblasts are responsible for the structural integrity of the extracellular matrix (ECM), its nutrient supply, and contribute to wound healing. However, CAFs present a tumor-like phenotype by engaging in cancer cell proliferation and invasion, angiogenesis, inflammation, immune system modulation, and ECM remodeling [10]. Healthy fibroblasts appear to modify their phenotypic profile to adapt to their new environment and progressively amplify the disease’s disastrous characteristics by shifting from passive responders to key effectors.

A variety of assumptions support the origin of CAFs activation and transformation, the most recent suggesting a role for metabolic alterations [12]. Indeed, various hallmarks of CAFs could be driven by a response to a wide range of paracrine signals (e.g., cytokines, chemokines, growth factors) emitted by cancer cells into the TME. The latter would induce CAFs to reprogram their energy production to aerobic glycolysis and produce high levels of energy-rich fuels for cancer cells to convert into biomass and proliferate faster in return. This process is known as the reverse Warburg effect [13-14].

An accurate mapping of all the knowledge related to CAFs involvement in the initiation and progression of cancer at the signaling, genetic, and metabolic levels would be beneficial to decipher the complex mechanisms underlying their activation. As knowledge assembly is an active field in systems biology, efforts have already been initiated with the publication of the CAF-map within the Atlas of Cancer Signaling Network (ACSN) [15]. The ACSN is a web-based multi-scale resource of biological maps depicting molecular processes in cancer cells and TME. It was developed in response to an increasing need for visualization and simplification of complex information resulting from massive production of biological data through high-throughput sequencing. The CAF-map is based on the manual curation of cell-specific molecular mechanisms from scientific literature and external databases. However, CAFs are a heterogeneous population within the TME [16] and their distribution across distinct cancer types is still under investigation. Despite its extensive coverage of the role of CAFs, the CAF-map, even updated in a cancer-specific manner, is limited regarding in-silico simulations by its static knowledge base function.

Nevertheless, it can be used as a basis to generate a cancer-specific CAF dynamic model to extend knowledge with executable information. Indeed, contributions of dynamic approaches to decipher metabolic reprogramming of fibroblasts have been highlighted recently to elucidate pathogenic mechanisms and accelerate identification of novel therapeutic targets [12].

Computational modeling is essential to gain insight on the emergent behavior of biological entities when complex and interconnected pathways are involved. Qualitative models based on logical relationships between components provide an appropriate description for systems whose mechanistic processes are unknown or lack quantitative data [17-18]. They allow a parameter-free study of the underlying dynamical properties of large biological pathways. Boolean models are the simplest form of qualitative models, yet extremely powerful [19]. They assume that all biological components are described by binary values (ON/OFF or 1/0) and their interactions by Boolean rules (“AND”, “OR”, “NOT”). Therefore, this formalism is very suitable for modeling signaling or gene regulation mechanisms carrying signal flow. In contrast, quantitative models provide great insight into biological systems by capturing their kinetic parameters [20]. For instance, constraint-based modeling allows fast calculations of large networks under steady-state assumptions [21]. This formalism is very appropriate to assess metabolic properties. Evidently, a hybrid formalism able to cover these various layers at once would be widely beneficial to study complex diseases such as cancer. Previous attempts in this direction include the PROM method, integrating high-throughput data into constraint-based modeling [22], the FlexFlux tool, combining flux balance analysis (FBA) and qualitative synchronous simulations [23], or the regulatory dynamic enzyme-cost flux balance analysis method (r-deFBA) combining metabolic dynamic modeling and transcriptional regulation [24]. Additionally, we recently developed a framework successfully generating a hybrid computational model of rheumatoid arthritis synovial fibroblasts, covering cellular signaling, gene regulation, and metabolism by coupling a cell- and disease-specific Boolean regulatory model with a constraint-based generic metabolic network [25]. Our approach improves on previous attempts by extracting information from asynchronous trap-spaces rather than synchronous stable states, providing a more comprehensive analysis of the regulatory model. Additionally, the previous need for user-defined transcription factors to target gene relationships or initial values and qualitative states to continuous intervals equivalences, a daunting task in large-scale regulatory models, is not required in our framework.

Computational modeling approaches have already been suggested to improve understanding of cancer notably with models of local invasion and migration [26], drug transport and delivery [27], or response to treatment [28]. The relevance of hybrid modeling approaches has also been recognized with the development of models integrating discrete and continuous variables to study tumor development and treatment [29]. However, regardless of the formalism (i.e., qualitative, quantitative, or hybrid), modeling efforts have not yet assessed the role of CAFs in the TME, let alone in a cancer-specific manner.

As breast cancer (BC) has overtaken lung cancer as the world’s most frequently diagnosed cancer since 2020 [11], the study of CAFs in this specific cancer has been initiated and some biological data are available.

To unravel the pathogenic mechanisms of cancer and accelerate the identification of innovative therapeutic targets, we aim to leverage our previously published coupling framework [25] and apply it to the integrative study of breast CAFs in the TME. Built upon previous efforts to formalize CAF-related knowledge in a detailed representation of disease mechanisms, we provide a state-of-the-art knowledge base further translated into a cell- and disease-specific Boolean model. By combining the latter regulatory model with a generic constraint-based metabolic network, this work presents the first large-scale hybrid model for breast CAFs. It addresses gene regulation, cellular signaling, and metabolic machineries in breast CAFs along with their involvement in the initiation and maintenance of the hallmarks of BC.

## 2 Methods

### 2.1 Update of the original CAF-map

The ACSN is an online database of multi-scale biological maps depicting molecular processes in cancer cells and TME [15]. All molecular interaction maps can be downloaded in various formats for further exploration and modifications. For this work, the CAF-map was downloaded from the ACSN 2.0 website^1^ in the standard XML format. It represents the major molecular interactions depicting the role of CAFs in the TME. However, the CAF-map displays generic CAF-specific interactions and is not specific to a certain type of cancer. It includes 681 components connected through 581 reactions and is based on the manual curation of 358 peer-reviewed articles. It is consistent with Systems Biology Graphical Notation (SBGN) standards [30] and thoroughly annotated with pertinent information and references. The CAF-map was imported on CellDesigner (version 4.4.2), a structured diagram editor for drawing regulatory and biochemical networks [31]. The original CAF-map was updated on multiple levels, following precise community-driven guidelines to ensure findability, accessibility, interoperability, and reproducibility, known as FAIR principles [32]. The CAF-map’s layout was edited to shift from a round cell with functional modules (e.g., “integrin signaling pathways”, “motility”, “growth factors production”, “cytokines and chemokines production”, “core signaling”) to a compartmentalized map with biological compartments (i.e., extracellular space, cytosol, nucleus, mitochondria, endoplasmic reticulum, secreted compartment, and phenotypes). With the help of specialized literature and external pathway databases (e.g., KEGG PATHWAY [33], PANTHER [34]), clear attribution of species to each compartment was obtained including modification or addition of interactions when needed. This representation allows for a better illustration of signal flow from top (displaying extracellular ligands complexation with plasmic receptors), through the cytoplasm (signaling and metabolic machineries), nucleus (gene regulation), to the bottom (secreted molecules and phenotype activation). Additionally, phenotypes were deleted either to allow for a detailed description of the pathways of interest (i.e., “TCA”, “glycolysis”, and “ketone bodies degradation” phenotypes), when they lacked added value in CAFs involvement in TME and cancer development (i.e., “microtubule polymerization”, “actin polymerization”, and “septine polymerization” phenotypes), or when their only interactions were to be activated by another phenotype (i.e., “stress fibril formation” and “matrix effects” phenotypes). Semantics were updated to comply with SBGN Process Description (PD) standards [35] for graphical visualization when not fully respected. It included adorning macromolecules with their residue (e.g., “phosphorylated”, “acetylated”) or state of modification (i.e., “active”, “inactive”). Components were named in accordance with HUGO Gene Nomenclature Committee (HGNC) IDs [36] for regulatory components and BiGG IDs [37] for metabolic species when not already. Relevant annotations regarding components and reactions (e.g., PubMed IDs (PMID), HUGO identifiers, pathways of interest) were previously stored in the note section of CellDesigner. Consistent with MIRIAM (Minimum Information Required In The Annotation of Models) standards [38], such notes were retrieved and re-inserted in the dedicated MIRIAM section of CellDesigner with the qualifier “bqbiol: isDescribedBy”. This qualifier is commonly used to link a component or a reaction to the literature or data describing it (e.g., PMIDs, DOI, GEO, KEGG identifier). Finally, an update of entities and reactions was carried based on the latest published mechanistic information and corrections or additions were applied if necessary. Only specific CAFs relevant studies were selected to enrich the map according to Curation and Annotation of Logical Models (CALM) standards [39].

The state-of-the-art and standardized CAF-map V2 was made available as an online interactive map on the standalone web server MINERVA (Molecular Interaction NEtwoRks VisuAlization) [40]. This platform allows for visual exploration, analysis, and management of large-scale molecular networks encoded in systems biology formats (e.g., CellDesigner, SBML, and SBGN). MINERVA also provides automated content annotation and verification along with overlaying experimental data.

### 2.2 Boolean model inference from the CAF-map V2

We used the CaSQ tool (version 1.1.0) [41] and its default parameters to generate a Boolean model from the CAF-map V2 in the standard Systems Biology Marked up Language-qualitative (SBML-*qual*) format [42]. Briefly, this map-to-model framework infers a Boolean model based on the topology and semantics of the molecular interaction map. Note that CaSQ allows to retain references, annotations, and layout of the molecular map in the associated model, facilitating interoperability.

The CAF-model was further made available on the Cell Collective repository of biological models [43].

### 2.3 Breast cancer contextualization of the CAF-model

As outlined above, the original CAF-map from the ACSN [15] was designed as a knowledge base comprising generic CAF-specific molecular mechanisms. Despite its updates and corrections, the CAF-map V2 remains a global map, gathering information from multiple types of cancer. This is consequently true for the Boolean model inferred from the latter.

Thus, a data-driven cancer-specific contextualization of the CAF-model was conducted. To this purpose, the dataset EGAD00001003808 from the European Genome-phenome Archive [44] was analyzed. It consists of RNA-Seq data from 47 CAF samples sorted from fresh BC including 28 CAFs-S1 and 19 CAFs-S4 subtypes. Both CAFs-S1 and CAFs-S4 subpopulations are myofibroblastic activated CAFs whose expressions are strictly restricted to cancer and characterized by high levels of fibroblast activating protein [45]. However, CAFs-S1 are mainly involved in tumor-like phenotypes (e.g., immunosuppression, tumor growth and proliferation, inflammation, ECM remodeling), whereas CAFs-S4 are responsible for generic core signaling, motility, and perivascular signatures. Thus, CAFs-S4 represented the control group against the aggressive CAFs-S1 group of interest. The tissue harvesting protocol, as well as the data mapping, alignment, quality control, and normalization processes are detailed in [10]. Differential expression analysis (DEA) was performed afterwards using the Limma package [46] in R (version 4.2.2) to identify differentially expressed genes (DEG) between breast CAFs-S1 and CAFs-S4. Standard significance threshold of adjusted p-value > 0.05 and absolute fold change (FC) > 1.5 were applied. Most significant DEG were depicted with an absolute FC > 1. Biological processes related to the most significant DEG were assessed by gene ontology enrichment analysis on the Gene Ontology Resource powered by PANTHER [47]. Identified DEGs were discretized to fix the CAF-model’s initial conditions in accordance with BC-specific biology. Up-regulated DEG present in the regulatory model were set to an initial value of 1. Similarly, down-regulated DEG present in the regulatory model were set to an initial value of 0. Finally, the inputs of a logical model being the nodes which do not present any upstream regulators, they are suspected to exert a significant control on the model’s dynamics due to the linearity of signal transduction. Thus, fixing their initial values is crucial to reproduce breast CAF-specific conditions. For inputs that would not have been fixed by the DEA, initial conditions may be extracted from BC-specific peer-reviewed literature. Overall, this effort to combine data-driven and manually curated breast CAF-specific information to initialize the regulatory CAF-model allows to confidently contextualize it to reproduce, as closely as possible, breast CAF-specific conditions.

### 2.4 Breast CAF-model regulatory behavior validation

Once contextualized, the breast CAF-model’s dynamic behavior was thoroughly assessed to confirm its biological relevance through two different approaches. First, a generic validation approach was carried to confirm the involvement of individual components into breast CAF-specific signaling or gene regulatory pathways. In greater detail, a literature review was undertaken and experimental evidence for breast CAF-specific activity were retrieved from in-vitro and in-vivo studies in both humans and murine models of BC (e.g., single or multiple knock-out and knock-in experiments, genetic recombination experiments). The latter studies experimental conditions were used to initialize the breast CAF-model (e.g., a gene knock-out experiment was translated as an initial value of 0 for said gene in the CAF-model and a gene knock-in was translated as an initial value of 1). This mechanistic verification was performed in a synthetic state of the model where remaining nodes were set to 0 (i.e., absent/inactive). Specific components to test values were then switched from 0 to 1 or 1 to 0 depending on the experimental scenario to assess their particular contributions to the pathway. Simulations were performed on the Cell Collective interactive platform [43] in the asynchronous updating mode, with a simulation speed of one, and a sliding window of 30. Secondly, the global behavior of the model was assessed. This comprehensive analysis, conducted at the level of the CAF-model’s cellular phenotypes (e.g., “growth factor production”, “immune system modulation”, “hypoxia”, “angiogenesis”, “cytokine production”) allowed to compare the model’s overall behavior to biologically known breast CAFs cellular behavior. This time, the model’s long-term behavior was evaluated through analysis of its trap-spaces under breast CAF-specific initial conditions identified earlier. Trap-spaces are regions of the state-space from which the system cannot escape. By definition, each trap-space contains at least one attractor of the model. However, trap-spaces may overlap with each other. Thus, minimal trap-spaces (later referred to as “trap-spaces”), or trap-spaces which do not include smaller trap-spaces, offer a good approximation of a model’s attractors, and faithfully capture the asymptotic behavior of Boolean models [48-49]. Trap-spaces were computed using BioLQM [50] directly using binary decision diagrams or an ASP-based solver. Their computation relies on the identification of positive and negative prime implicants for each component’s function without performing simulation but rather through a symbolic approach implementing a constraint-solving method. In this context, each trap-space reflects a different subspace of breast CAFs cellular phenotypes. The latter were compared with known breast CAF overall cellular behavior extracted from the literature to confirm the model’s asymptotic behavior.

### 2.5 Breast CAF-model coupling with a generic metabolic model

The breast CAF-model was further integrated with a generic core metabolic network, MitoCore [51], to cover an additional biological layer of metabolism. MitoCore is a manually curated constraint-based model of human central metabolism, including 74 metabolites, 324 metabolic reactions, and 83 transport reactions. Although initially parameterized for human cardiomyocytes, MitoCore’s default parameters can be extended to various biological contexts without implying cell-specific features. Its realistic ATP production rates for different substrates have been proven [51], making it very valuable in the present study as opposed to larger genome-scale reconstruction of human metabolism exhibiting unrealistic results.

The coupling of the regulatory breast CAF-model with the generic MitoCore metabolic network was conducted by exploiting the asymptotic behavior of metabolic components present in both models to extract additional regulatory-based metabolic constraints. In particular, metabolic components exhibiting an asymptotic inactive behavior under regulatory influences (**Figure 1**).

**Figure 1.**
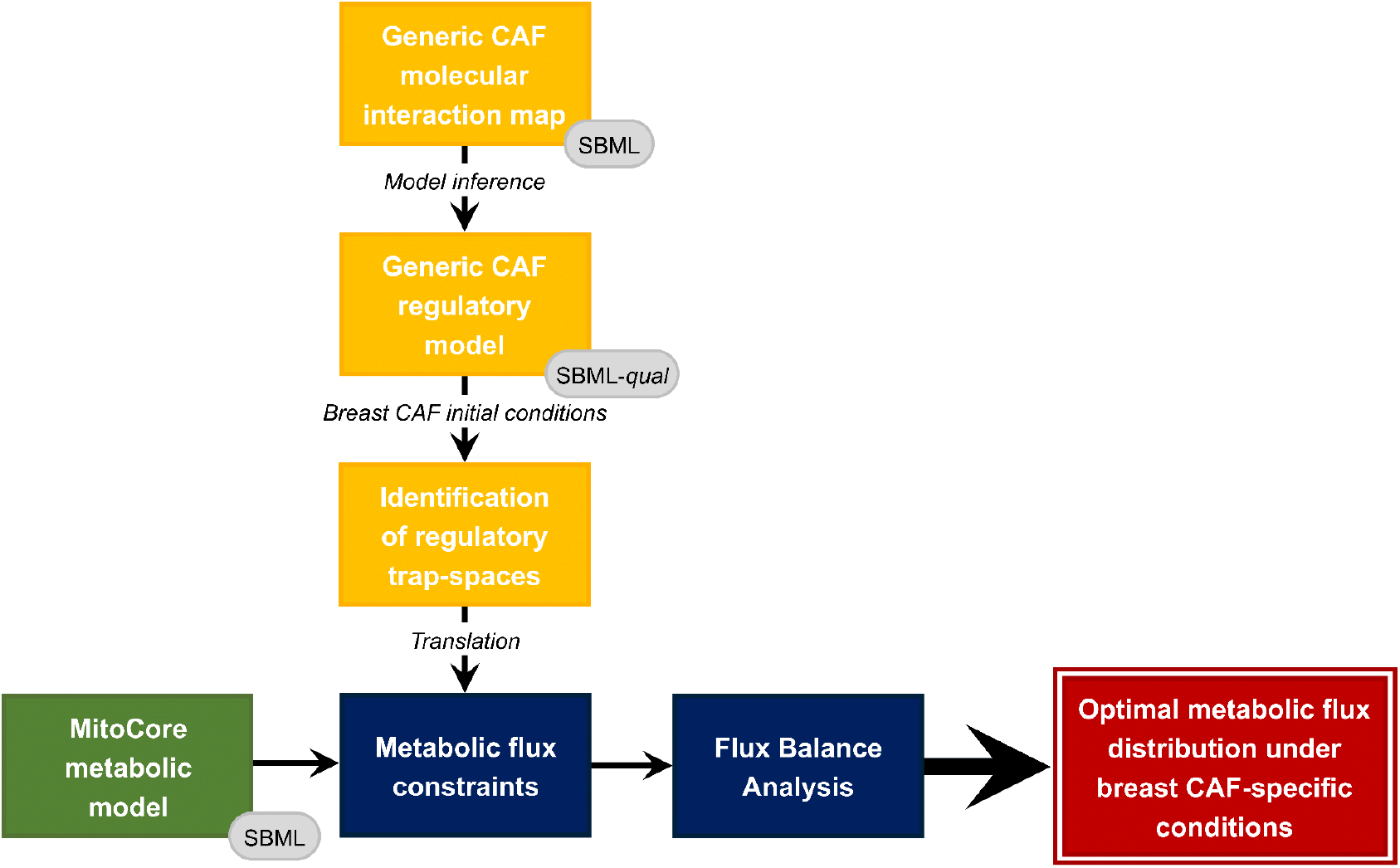
General architecture of the hybrid modeling framework.

Briefly, cell- and disease-specific initial conditions were used to initialize the regulatory CAF-model before applying the value propagation algorithm [52] implemented in CoLoMoTo [53]. The latter allowed to evaluate the dynamical influence of initial conditions on the global behavior of the model to decrease its complexity. In this restricted space, minimal trap-spaces of the regulatory network were identified, reflecting its asymptotic behavior, and particularly the asymptotic behavior of metabolic components (i.e., metabolic enzymes and metabolites) influenced by cell- and disease-specific signaling and gene regulatory processes. The maximal value of trap-spaces relative to metabolic components were used to constrain associated MitoCore’s metabolic fluxes. For every metabolic component with a projected maximal regulatory trap-space equal to 0, the flux of its associated MitoCore metabolic reactions (i.e., catalyzed reactions for enzymes or producing reactions for metabolites) were constrained to 0. This approach assumes that a metabolic enzyme with a projected maximal trap-space value equal to 0 (i.e., trap-spaces always equal to 0) expresses the inactivation of the associated enzyme by signaling or gene regulation pathways. The reactions catalyzed by the enzyme will not happen. Similarly, a metabolite-associated maximal trap-space value of 0 denotes the absence of production of the associated metabolite by signaling or gene regulation pathways. Its producing reactions did not occur. Alternatively, metabolic-associated trap-space values different than 0 would reflect a potential activation or production of the associated species. This qualitative information does not reflect the feasibility nor the kinetics of the associated reactions. It is not sufficient to extract additional metabolic constraints. By extracting additional constraints from metabolic components with a proven asymptotic inactive behavior under cell- and disease-specific regulatory conditions, our framework enables contextualization of MitoCore’s generic metabolic model in a breast CAF-specific manner. For further details, the general architecture of the framework for hybrid modeling is provided and thoroughly detailed in [24].

### 2.6 Metabolic network analysis

A widely used method for analyzing large-scale reconstructions of metabolic networks is FBA [54]. Its main advantage lies in the need for little information regarding the kinetic parameters of biological components as it calculates metabolic fluxes by assuming steady state conditions. Two FBAs were performed using the python package CobraPy [55] to highlight a potential change in fibroblast metabolism under disease-specific conditions. First FBA was conducted without any additional constraints from the regulatory network and considered as the “control”, reflecting the healthy fibroblast state. Second FBA was conducted with additional metabolic constraints extracted from the breast CAF-specific regulatory model’s trap-spaces, reflecting breast CAFs condition. The FBA objective function was set to maximal cellular energy production in the form of ATP to reflect the central role of metabolism. The latter was manually defined as the sum of the three ATP producing reactions, i.e., step seven and ten of glycolysis and complex V of oxidative phosphorylation (OXPHOS). As MitoCore lacks cellular and tissular specificity, a numerical interpretation of metabolic flux values is not possible. FBA results were rather interpreted in terms of flux distribution through the ratios of ATP production from glycolytic (i.e., fluxes of step seven and ten of glycolysis) or oxidative reactions (i.e., flux of OXPHOS complex V reaction) relative to total ATP production (i.e., objective function flux) to highlight the main metabolic pathway of ATP production. Carbon fluxes (C-flux) of uptake and secretion were also analyzed to investigate the exchanges between the cell (i.e., breast CAF) and its environment (i.e., breast TME). They respectively represent the total cellular carbon influx from a specific uptake reaction and the proportion of total cellular carbon efflux from a specific secretion reaction. Finally, a comparison of internal metabolic fluxes is performed, leading to identify metabolic alterations from a greater than 2-fold variation.

### 2.7 Identification of regulatory molecular drivers

Beyond the overall impact of breast CAFs cellular signaling and gene regulatory pathways upon their metabolic machinery, we aim to identify key regulators. As stated above, the regulatory network’s inputs are suspected to significantly influence the model’s dynamics due to the linearity of signal flow. Hence, successive individual knock-outs and knock-ins of regulatory inputs were conducted to identify potential key regulators. All regulatory inputs initially fixed at 1 under breast CAF-specific initial conditions were successively fixed at 0 while the other components remained at breast CAF-specific values, mimicking experimental knock-outs. Similarly, all regulatory inputs fixed at 0 under breast CAF-specific initial conditions were successively fixed at 1 while the others remained unchanged, mimicking experimental knock-ins. Each new set of initial conditions was used to initialize the CAF-model and the framework for hybrid modeling was applied as outlined above. Each FBA result illustrates the distribution of metabolic fluxes under new regulatory influences. Again, the proportion of total energy production in the form of cellular ATP production from glycolytic and oxidative pathways was reported to identify a potential alteration under new regulatory conditions.

## 3 Results

### 3.1 CAF-map V2

The CAF-map V2 is available on the MINERVA platform^2^. This extensive knowledge base illustrates the major cellular signaling, gene regulation, and metabolic pathways as well as molecular mechanisms and phenotypes involving CAFs in cancer initiation and progression.

It is fully compliant with SBGN PD standards for visualization [35], MIRIAM for annotation [38], and CALM for bio-curation [39]. It includes 649 species (308 proteins, 95 genes, 114 RNAs, 43 simple molecules, and 89 molecular complexes), 19 phenotypes, and 544 reactions. A dozen additional peer-reviewed articles enabled corrections and extended the mechanical processes depicted in the map for a total of 368 PMIDs. This high number of references enhances confidence in presented mechanisms. Biological compartments depicting extracellular space, nucleus, mitochondrion, endoplasmic reticulum, secreted compartments, and phenotypes were further added in the map to account for biological compartmentalization and cell transport. Beyond the considerable effort of the initial CAF-map to include major cancer functional modules (e.g., “growth factors production”, “cytokines”, “chemokines production”, “matrix regulation”) and molecular pathways (e.g., “growth factors signaling pathways”, “inflammatory signaling pathways”, “integrin signaling pathways”), coverage in disease-specific mechanisms was expanded in the CAF-map V2. For instance, metabolic pathways initially only considered as phenotypes were fully detailed, along with addition of the calcium ion signaling pathway in the new endoplasmic reticulum compartment. Visualizations of the original CAF-map and CAF-map V2 are provided in **Figure 2**.

**Figure 2.**
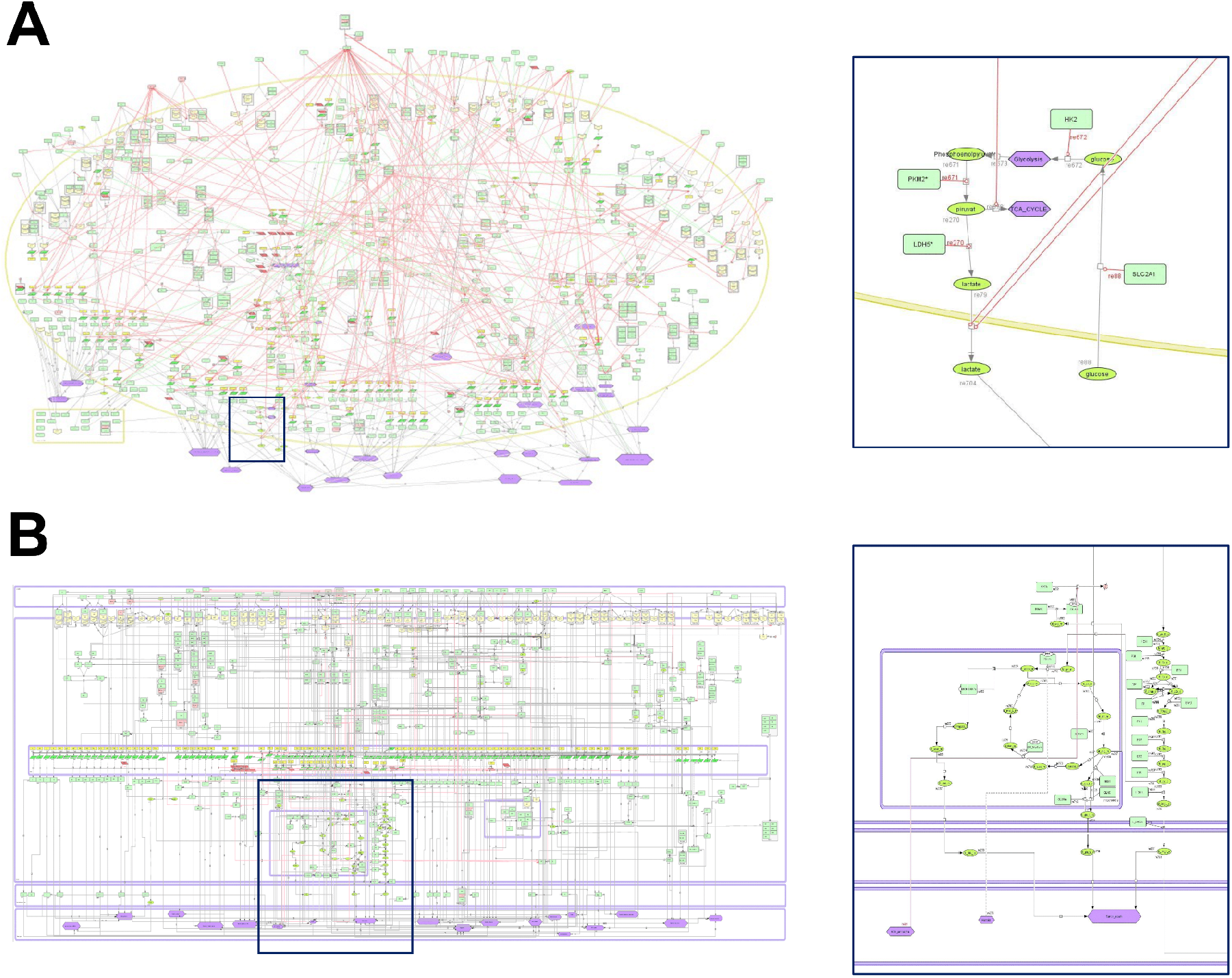
CellDesigner visualization of the **(A)** original CAF-map from ACSN [14] and **(B)** CAF-map V2 with associated zoom-ins on glucose-related pathways. The CAF-map V2 depicts a standardized formal representation and layout revision with a clear signal flow. Components are precisely attributed to biologically relevant compartments. Glucose pathways are compartmentalized and are no longer represented only as phenotypes but properly detailed. Up-to-date mechanistic information was further added.

### 3.2 Breast CAF-model

Translation of the CAF-map V2 using CaSQ generated a dynamic Boolean model of 463 nodes (including 62 inputs) and 793 interactions. The latter is publicly available on the Cell Collective platform^3^. However, components and interactions depicted in the model are not cancer-specific but generic to CAFs.

The DEA conducted to contextualize the model identified 3678 DEG in CAFs-S1 vs. CAFs-S4 among the 18252 mapped genes. 1866 DEG were significantly up-regulated in CAFs-S1 vs. CAFs-S4, leading to fix an initial condition of 1 for 71 nodes of the CAF-model. Most significantly up-regulated DEG in CAFs-S1 are mainly involved in chemotaxis, ECM organization, locomotion, response to growth factors, cell adhesion, migration, motility, differentiation, and proliferation. Such observations support the classification of CAFs-S1 as the subpopulation of CAFs carrying key aggressive functions. 1812 DEG were significantly down-regulated in CAFs-S1 vs. CAFs-S4, leading to fix an initial condition of 0 for 34 nodes of the CAF-model. Most significantly down-regulated DEG in CAFs-S1 are involved in ion transport and signaling, specifically calcium ions. Finally, 41 CAF-model’s inputs remained unfixed, leading to assign their values through manual curation of peer-reviewed breast CAF-specific literature. The complete list of breast CAF-specific initial conditions along with their source of attribution is outlined in **Table S1** of supplementary material.

To validate the behavior of the model, generic model simulations were first compared to breast CAF-specific experimental scenarios extracted from the literature. Details of each experimental scenario, associated CAF-model initialization, and dynamic results are presented in **Table S2** of the supplementary materials. Overall, regarding this generic evaluation, 29 experimental scenarios were confirmed by the CAF-model out of 41 biological scenarios. Ten scenarios were not reproducible mostly due to a lack of mechanistic detail regarding specific interactions in the literature, leading to a missing or incomplete representation in the CAF-map V2 and the associated breast CAF-model. Finally, two generic scenarios were not validated as other pathways were needed concomitantly. Subsequent global model’s behavior evaluation under breast CAF-specific initial conditions, i.e., identification of its complete asymptotic behavior, depicted 128 trap-spaces (see **Table S3** of the supplementary materials). Each trap-space reflects a different subspace of breast CAFs cellular phenotypes. Most components values are stable within all trap-spaces (always fixed at 0 or 1) but others vary within trap-spaces. The projection of trap-spaces restricted to the model’s ontological phenotypes (i.e., distinct cellular outcome) identifies a single trap-space (**Table 1**). The latter illustrates activity of aggressive phenotypes in breast CAFs, e.g., angiogenesis, fibroblast proliferation, hypoxia, matrix degradation, tumor growth and invasion, while negative regulators are inactive. These findings are consistent with the behavior of breast CAFs as described in scientific literature [56].

**Table 1.**
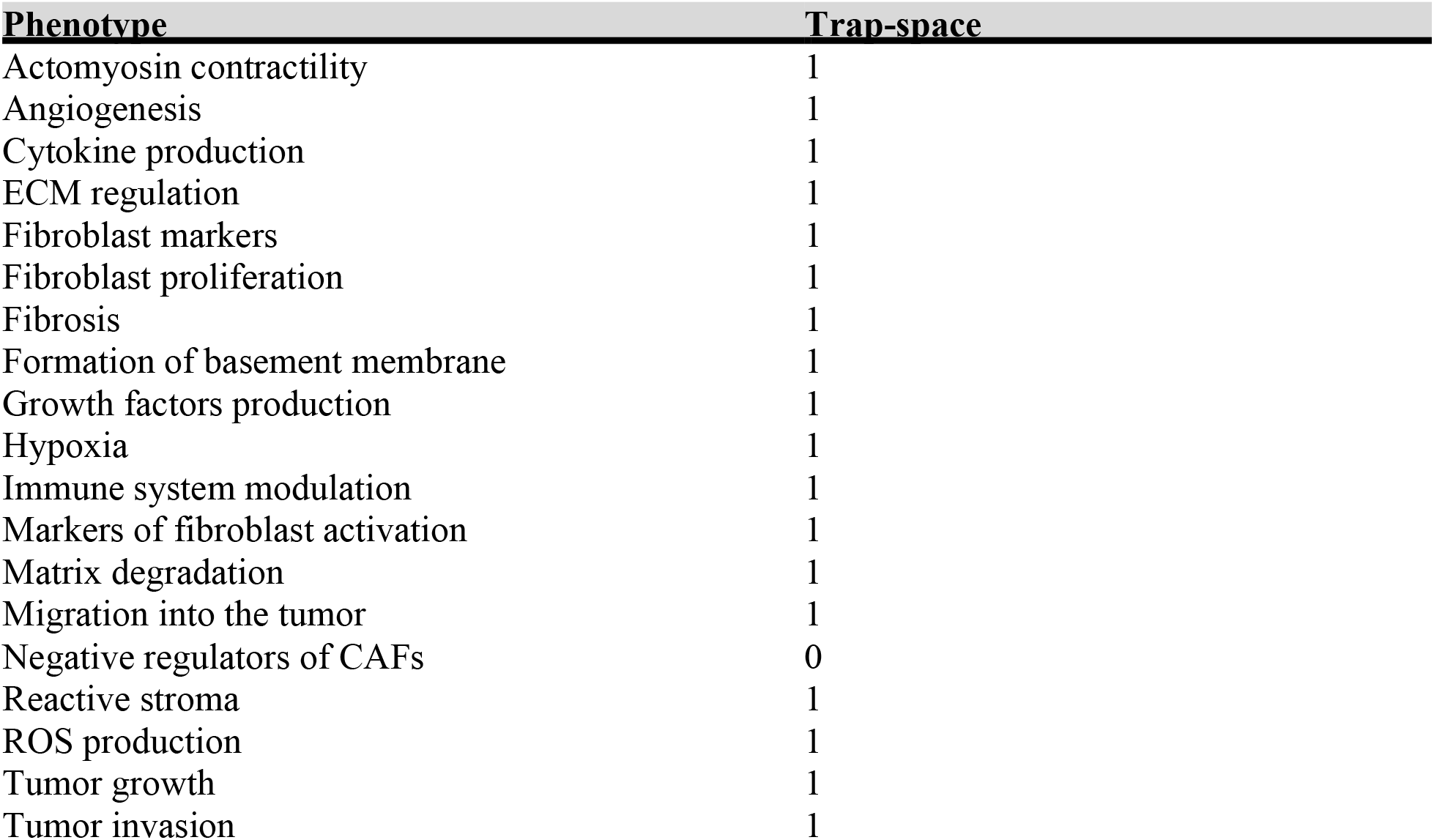
Projection of the breast CAF-model’s trap-spaces restricted to its ontological phenotypes.

The asymptotic state of ontological phenotypes is the result of their regulators combined effect exerted through their Boolean rules. Thus, the behavior of groups of biomarkers associated with each ontological phenotype can be identified to further validate the behavior of the breast CAF-model. For instance, the constant activation of matrix metalloproteinases (e.g., MMP2, MMP9), known to lead to matrix degradation [57], within all trap-spaces, accounts for the asymptotic activation of the associated phenotype. Their role in the coordination of ECM is also well established [56], which supports the consistency of their trap-space with the “ECM regulation” phenotype. Likewise, most interleukins (e.g., IL1A, IL1B, IL11 IL6) being active under breast CAF-specific conditions confirms their experimentally observed function of immune system modulation and cytokine production drivers [58]. The asymptotic activated state of the hypoxic phenotype is explained by the sustained activation of its main molecular driver, Hypoxia -Inducible Factor 1 (HIF-1) [59]. The latter is further involved with ROS production [59], confirmed in the trap-spaces analysis. Likewise with the angiogenic and reactive stroma phenotypes and their key player, VEGF [60]. Migration, tumor growth, and invasion appear to be driven by growth factors and Wnt signaling pathway, consistent with literature [61]. Furthermore, activation of key fibroblastic markers (e.g., FAP, PDGF) in parallel with inactivation of known negative regulators of breast CAFs (e.g., miR101, miR141, CAV3, SIRT3) under initial disease-specific conditions explain the state of both related phenotypes. Finally, fibroblast proliferation requires EGF and TGFB signaling pathways [62]. Fibrosis and basement membrane formation seems governed by collagens [63]. Actin, myosin, and TNC drive actomyosin contractility [64], all confirming experimental observations in breast CAFs.

### 3.3 Hybrid breast CAF-model

When projected on the breast CAF-model’s metabolic components, i.e., metabolic enzymes and metabolites, a single trap-space is identified. It is a stable state, which indicates that, under influences from breast CAF-specific cellular signaling and gene regulation networks, the asymptotic behavior of all metabolic components is fully stable. As described in the hybrid modeling framework, additional metabolic constraints were extracted from metabolic compounds with a proven inactive asymptotic behavior. This concerns 7 metabolic enzymes and 15 metabolites leading to constrain the flux of 66 unique metabolic reactions to 0 (see **Table S4** and **Table S5** of supplementary materials). The high number of constrained reactions is explained by enzymes commonly catalyzing a single metabolic reaction whereas metabolites were produced by numerous reactions. Additionally, certain metabolic reactions (e.g., R_PDHm, R_ICDHxm, R_GLUNm) were constrained both through their catalyzing enzymes and their produced metabolites. It reflects the consistency of breast CAF-specific cellular signaling and gene regulation machinery upon its metabolic processes. This approach allows to contextualize MitoCore, a generic core metabolic network, in the cell- and disease-specific context of breast CAFs.

### 3.4 Metabolic network analysis

Visualization of both control and breast CAF-specific FBA results are provided in **Figure 3A** and **3B**, details of metabolic fluxes in both conditions are provided in **Table S6** and **Table S7** of supplementary materials. Optimal fluxes for ATP production in a healthy fibroblast are OXPHOS fluxes, accounting for 96% of cellular ATP production. Main uptaken carbonated molecules are hexadecanoate (40.92% of total C-flux), glucose (29.48%), lactate (9.42%), and HCO_3_ (9.34%), primary energy sources for most cells. Main secreted carbonated molecule is CO_2_ (99.96%). Under breast CAF-specific regulatory conditions, optimal fluxes for ATP production are glycolytic fluxes, as they now explain 85.05% of cellular energy production in the form of ATP. Main uptaken carbonated molecules are glucose (87.72% of total C-flux) and aspartate (10.01%). Main secreted carbonated molecules are lactate (88.69%) and alanine (7.99%). Comparison of internal metabolic fluxes revealed reprogramming of major metabolic pathways for ATP production. Globally increased glycolytic fluxes along with increased lactate secretion in breast CAFs reflect a highly glycolytic metabolism. Low oxidative metabolism is demonstrated by decreased OXPHOS and TCA-associated fluxes along with decreased secretion of oxidative by-products (e.g., CO_2_ and H_2_O). A hypoxic environment along with anaerobic metabolism is revealed through decreased O_2_ uptake and increased H^+^ secretion, associated with environmental acidity. Besides metabolic pathways for energy production, macromolecular building blocks pathways are altered in breast CAFs. Particularly, amino-acids uptake by breast CAFs is decreased (e.g., proline, glycine), along with their decreased degradation (e.g., proline), and increased secretion (e.g., serine, lysine). Fatty acids uptake is further decreased (e.g., butanoic acid) while their secretion is increased (e.g., palmitic acid). Cardiolipin synthesis is decreased in breast CAFs, coherently with former findings as they are known to regulate OXPHOS. Further pathways including folate cytosolic, reductive carboxylation, and butanoate metabolism, appear to be impacted, most likely resulting indirectly from metabolites redirection through other altered pathways or through the application of our framework’s additional metabolic constraints. Finally, mitochondrial transporters are affected due to the reprogramming of mitochondrial pathways discussed above.

**Figure 3.**
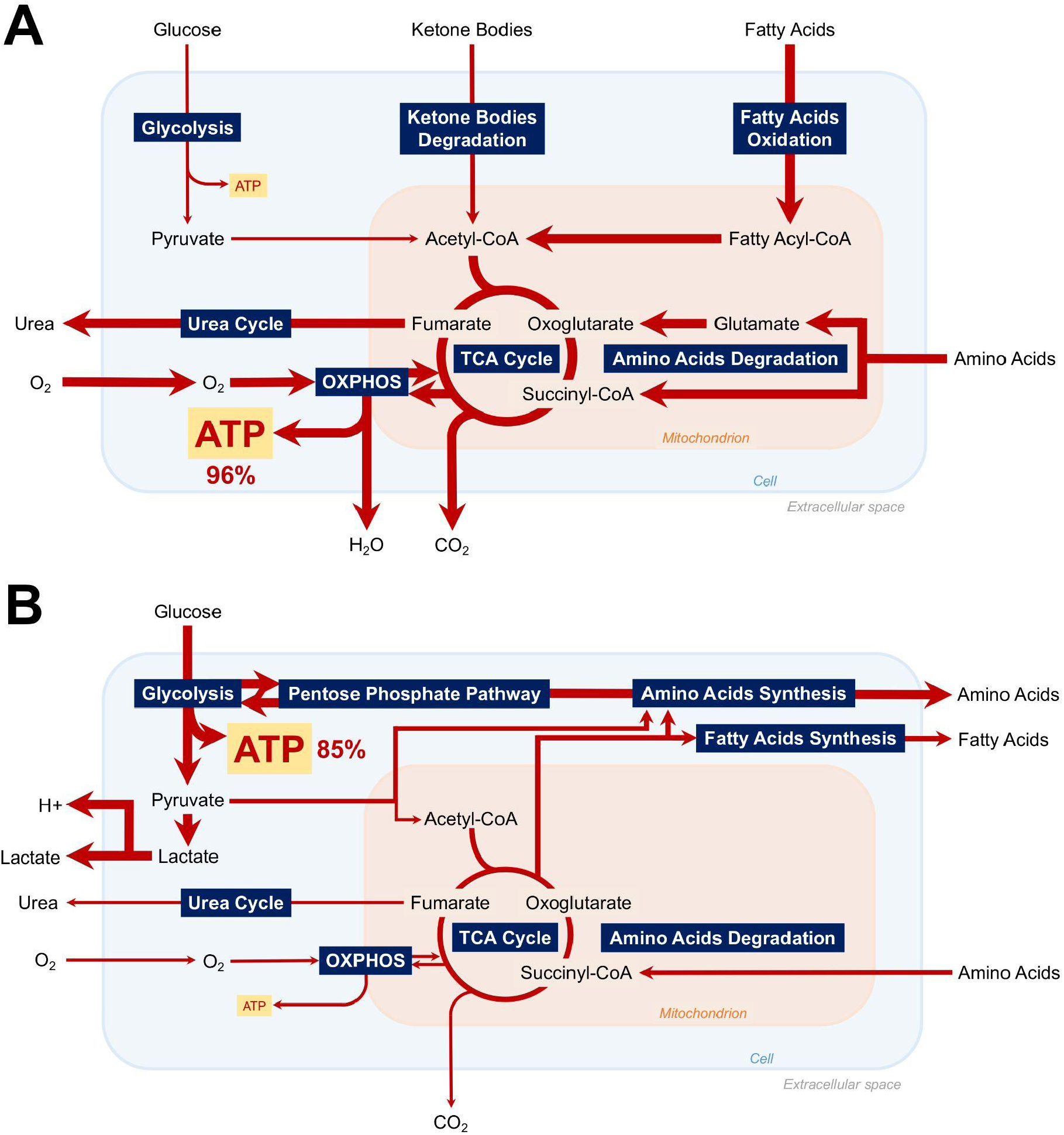
Visualization of major active metabolic pathways for maximal cellular ATP production in **(A)** control and **(B)** breast CAF-specific conditions according to Flux Balance Analysis.

### 3.5 HIF-1, regulatory driver of metabolic reprogramming in breast CAFs

After computationally reproducing the experimentally observed metabolic reprogramming in breast CAFs, the challenge consists in identifying its main regulatory drivers. Successive individual knock-outs and knock-ins of breast-CAF specific regulatory inputs resulted in 147 new sets of initial conditions, and therefore 147 new FBAs (complete list of FBA results per set of initial conditions by ratio of total cellular ATP produced through glycolytic or oxidative pathways is provided in **Table S8** of supplementary materials). Among the performed FBA, only one set of regulatory initial conditions reproduces a healthy metabolic profile for energy production in breast CAFs, it is condition 57 (C57). Indeed, by knocking-out HIF-1, metabolic pathways seem to recover a healthy-like distribution with cellular ATP being generated from oxidative rather than glycolytic pathways. Other regulatory inputs do not seem to directly affect metabolic fluxes distribution in breast CAFs.

## 4 Discussion

To investigate the complex mechanisms underlying the key role of CAFs in TME for breast cancer development and maintenance, dynamic modeling is essential. In this regard, we developed the first large-scale Boolean model for breast CAFs. It covers signaling, gene regulation, and their impact upon metabolism in a cell- and disease-specific context. As a basis, the CAF-map from the ASCN resource [15] was updated to fully meet community-driven guidelines for representation [30, 35], curation [39], and annotation [38] of biological data. This extensive manually constructed knowledge base provided the basis to infer a Boolean model through CaSQ tool’s map-to-model framework [41]. The CAF-model was further contextualized to reproduce breast CAF-specific conditions. It was achieved by combining both data-driven and manual curation approaches to ensure high cellular specificity and confidence in the depicted interactions and initial conditions. Biological coherence of the breast CAF-model’s behavior was assessed against experimental evidence extracted from the scientific literature at two distinct levels (i.e., generic pathways and global cellular behavior), each time reproducing experimental observations. Overall, generating and parameterizing a logical model from manual network construction along with data analysis and bio-curation allows to avoid issues associated with automatic and non-curated methodologies (e.g., improper reconstruction of reactions, incorrect compartmentalization or representation, lack of omics data in poorly addressed biological fields).

As displayed in our previously published workflow [25], integration of the breast CAF-specific regulatory model with MitoCore’s generic reconstruction of human central metabolism [51] in a context-specific manner was achieved through extraction of additional constraints from metabolic components with a proven inactive state. We further demonstrated the adaptability of our hybrid modeling framework as it successfully coupled a new regulatory network with MitoCore. Built upon previous attempts to couple regulatory with metabolic networks [22-24], we proposed integrating a core metabolic network through interpretation of regulatory trap-spaces. It allows the assessment of the impact of its long-term regulatory behavior on the updating of metabolic sub-systems under cell- and disease-specific conditions. This approach, while capturing high-level of regulatory complexity, overcomes the need for translating discrete synchronous qualitative states into user-defined continuous intervals as well as relying solely on automated integration of high-throughput data. Overall, combining various layers of biological machinery, namely signaling, gene regulation, and metabolism, enabled a rich characterization of the role of CAFs in breast TME.

Simulations of our hybrid network constitute the first modeling attempt to address breast CAFs regulatory impact upon their metabolism. Findings depicted a hypoxic and acidic TME along with a highly glycolytic metabolism and almost null oxidative fluxes for energy production in breast CAFs. Increased production and secretion of energy-rich fuels (e.g., pyruvate, lactate, amino-acids, fatty acids) along with decreased secretion of oxidative by-products (e.g., CO_2_, H_2_O) were reported, fully reproducing the experimentally observed reverse Warburg effect in CAFs [65]. Fibroblasts would undergo metabolic reprogramming and turn into “metabolic slaves”, generating high levels of energy-rich fuels through pentose phosphate pathway or amino acids synthesis. These nutrients would feed cancer cells in macromolecular building blocks and sustain their aggressive phenotype. Such metabolic alterations are suspected to be driven directly by signals from cancer cells, illustrating an extensive crosstalk within the TME. Subsequent simulations identified HIF-1 as the regulatory activating stimuli of breast CAFs metabolic reprogramming. HIF-1 is known as a master transcriptional factor involved in homeostasis and cellular response to hypoxia. Indeed, chronic hypoxia is a major hallmark of solid tumors as they quickly outgrow their blood supply to support their continuous growth and proliferation, leaving parts of the tumor with almost null concentration of oxygen. Angiogenesis, metastasis, and drug resistance benefit from this hypoxic state [66]. To sustain such aggressive behaviors in this challenging environment, cancer cells need additional fueling. As a result, this chronic hypoxic and acidic environment, generated by cancer cells to maintain their tumorigenic phenotype, was recently suspected to induce breast CAFs reverse Warburg effect and promote BC progression [66]. Our findings fully support this hypothesis and further identify HIF-1 as a main molecular regulator. In response to cancer cells hypoxic paracrine signals, breast CAFs would activate transcription of glycolytic genes along with glucose transporters, suppress oxygen consumption, by-pass oxidative pathways, and induce a reverse Warburg effect. Thus, we hypothesize that targeting the metabolic reprogramming of fibroblasts through HIF-1’s pro-glycolytic and anti-oxidative transcriptional activity within a therapeutic strategy including the TME could benefit the treatment of BC. Similar in-silico experiments were previously conducted in [25] with a hybrid model of rheumatoid arthritis synovial fibroblasts (RASFs). We suggested a cancer-like reverse Warburg effect in the rheumatic joint where RASFs would reprogram their metabolism to produce more ATP, maintain their aggressive phenotype, and feed neighboring energetically demanding cells (e.g., chondrocytes, macrophages, dendritic cells) with fuels and nutrients. Metabolic reprogramming of fibroblasts appears to be a crucial element in the pathogenesis of two complex diseases as different as BC and rheumatoid arthritis (RA). Due to the similarity of the framework applied to contextualize the generic MitoCore network in both diseases, a comparison of altered metabolic pathways can be performed (see **Table S9** of supplementary materials for RASF-specific FBA results). Overall, metabolic pathways directly related to energy production are altered in the same manner (i.e., increased glycolysis, decreased OXPHOS and TCA). Oxidative by-products are accordingly altered in both breast CAFs and RASFs (e.g., increased lactate and secretion, decreased CO_2_ and H_2_O secretion). The fibroblasts environment, respectively BC TME and RA joint, appears to be similarly modified (e.g., decreased O_2_ uptake and increased H^+^ secretion). Metabolic pathways not directly involved in ATP production are additionally affected. For instance, disease-specific regulatory conditions similarly reprogram amino-acids and fatty acids pathways in RA and BC-associated fibroblasts probably due to their importance in the biosynthesis of macromolecules for aggressive cells. Indeed, secretion of building blocks is additionally increased in both situations. Certain pathways are similarly altered but raise the question of their interest under different environmental conditions. For instance, reductive carboxylation is similarly increased in breast CAFs and RASFs. Acting as a novel glutamine pathway, it supports the growth of cells depicting mitochondrial deficiencies. Warburg originally hypothesized that cancer-like cells presented a mitochondrial defect [68] but later work refuted it [69]. Such studies have not yet been conducted in both types of fibroblasts. Further experimental investigations are needed to decipher their mitochondrial status and identify a potential benefit from reductive carboxylation reactions. Same goes for cytosolic misc, needing additional experimental studies to decipher their precise beneficiaries in BC and RA. Altogether, mitochondrial and cytosolic transporters or shuttles pathways are affected, not necessarily in the same way, but all due to the reprogramming of upstream metabolic pathways producing their metabolite of interest. Finally, butanoate metabolism is not affected similarly in breast CAFs and RASFs. However, as this pathway is typically involved in processes associated with intestinal fermentation, its alterations do not seem to be significant in a cancerous or autoimmune context.

Beyond the many shared altered metabolic pathways, the key molecular regulator was also identified as HIF-1 in RASFs reverse Warburg effect where its mechanism of action was suspected at the transcriptional level. Already recognized in RA as a driver of inflammation, angiogenesis, and cartilage destruction [70], targeting HIF-1 has not yet been proposed within a therapeutic strategy aiming at the resolution of metabolic reprogramming in fibroblasts. In BC, therapeutic opportunities targeting HIF-1 appeared until very recently to be limited to its metastatic or driver of tumor-proliferation activity [71]. Growing interest in metabolic targeting to address pro-tumor characteristics resulted in a few insights such as Honokiol as an inhibitor of HIF-1-mediated glycolysis to halt BC cells growth [72]. However, its interest in breast CAFs has not yet been investigated. Finally, studies in other types of human fibroblasts have recognized a key role for HIF-1 (e.g., in anti-aging and regeneration in dermal fibroblasts [73], attenuating fibrosis and delaying vascular remodeling in systemic sclerosis-associated fibroblasts [74] but its targeting in the resolution of fibroblasts metabolic reprogramming has not yet been studied.

According to our findings and given the shared metabolic alterations and regulatory molecular driver, we assume the existence of a common mechanism directing the phenotypic transformation of fibroblasts through a HIF-1-driven metabolic reprogramming in BC and RA. In both situations, regulation of CAFs and RASFs through a reverse Warburg effect enables them to adapt and survive in a new hypoxic and acidic environment, maintain their surroundings, and actively participate in the amplification of the associated disease’s debilitating symptoms.

## 5 Conclusion

Hybrid modeling formalisms are highly relevant for the study of complex diseases such as cancer. By covering multiple levels of intertwined processes and biological entities, they contribute to deciphering the complex mechanisms at the origin of carcinogenesis and further develop therapeutic strategies considering tumor cells surroundings. In this regard, we present the first large-scale hybrid breast CAF-model covering major cellular signaling, gene regulation, and metabolic processes. It combines an asynchronous cell- and disease-specific Boolean model with a generic core metabolic network. Generation of this hybrid model leveraged both data-driven and manual curation approaches to ensure high cellular specificity and confidence in the depicted interactions and contextualization. Hybrid simulations reproduce the experimentally observed reverse Warburg effect in breast CAFs and further identify HIF-1 as its primary molecular driver. Following paracrine hypoxic signals emitted by cancer cells in the TME, breast CAFs would reprogram their energy production pathways and produce high-levels of energy-rich fuels and nutrients for neighboring tumor cells. Developing innovative treatment strategies targeting the reverse Warburg effect in CAFs through HIF-1 may represent a promising path for the treatment of breast cancer. This process appears to be further shared with rheumatoid arthritis synovial fibroblasts.

## 6 Perspectives

The presented approach, while capturing the complexity of signaling and gene regulation processes upon breast CAFs metabolism, provides a linear view of biological events. Considering the regulatory network’s update from metabolic outputs may provide a more comprehensive understanding of events associated with the reverse Warburg effect in breast CAFs. Additionally, identification of regulatory molecular drivers for breast CAFs metabolic reprogramming focuses solely on regulatory inputs. The effect of internal core components is not considered although it could enable identification of HIF-1’s precise mechanism of action. Finally, the regulatory inputs are tested individually but biological joint influences cannot be dismissed. Combined knock-outs and knock-ins could be considered, leading to therapeutic strategy suggestions targeting several components at once.

## Abbreviations

ACSN: atlas of cancer signaling network
BC: breast cancer
C-flux: carbon flux
CAFs: cancer-associated fibroblasts
CALM: curation and annotation of logical models
DEA: differential expression analysis
DEG: differentially expressed genes
ECM: extracellular matrix
FAIR: findability accessibility interoperability and reproducibility
FBA: flux balance analysis
FC: fold-change
HGNC: HUGO gene nomenclature committee
HIF-1: hypoxia-inducible factor 1
MINERVA: molecular interaction networks visualization
MIRIAM: minimum information required in the annotation of models
OXPHOS: oxidative phosphorylation
PD: process description
PMID: PubMed ID
RA: rheumatoid arthritis
RASFs: rheumatoid arthritis synovial fibroblasts
SBGN: systems biology graphical notation
SBML-*qual*: systems biology marked up language-qualitative
TCA: citric acid cycle
TME: tumor microenvironment.

## Acknowledgements

The authors would like to thank Marek Ostaszewski (Luxembourg Centre for Systems Biomedicine) for his technical help handling CellDesigner annotations and hosting the CAF-map V2 on MINERVA, as well as Fatima Mechta-Grigoriou and Yann Kieffer (Institut Curie) for granting access to the breast CAFs RNA-Seq data (European Genome-phenome Archive dataset EGAD00001003808).

## Authors Contribution

Conceptualization: Sylvain Soliman, Anna Niarakis; Data curation: Sahar Aghakhani, Sacha E Silva-Saffar; Formal analysis: Sahar Aghakhani, Sacha E Silva-Saffar; Funding acquisition: Anna Niarakis, Sylvain Soliman; Investigation: Sahar Aghakhani, Sacha E Silva-Saffar; Methodology: Sahar Aghakhani, Sylvain Soliman, Anna Niarakis; Project administration: Sylvain Soliman, Anna Niarakis; Resources: Sylvain Soliman, Anna Niarakis; Supervision: Sylvain Soliman, Anna Niarakis; Validation: Sahar Aghakhani, Sacha E Silva-Saffar, Sylvain Soliman; Visualization: Sahar Aghakhani; Writing — original draft: Sahar Aghakhani; Writing — review & editing: Sahar Aghakhani, Sylvain Soliman, Anna Niarakis.

## Footnotes

1 https://acsn.curie.fr/ACSN2/downloads.html

2 https://pathwaylab.elixir-luxembourg.org/minerva/index.xhtml?id=CAF-map_V2 (temporary link)

3 https://research.cellcollective.org/#30b2359b-38be-42cc-8885-c3ab9b8f8063 (temporary link)

