## Supplementary material for "Hybrid computational modeling highlights reverse Warburg effect in breast cancer-associated fibroblasts": Supplementary_materials_TABLE-S1.pdf

**Table S1. Breast CAFs-specific initial conditions generated from data-driven differential expression analysis (DEA) or manual curation of peer-reviewed literature (PubMed IDs are provided).**

| Component | Function | Initial value | Source |
|---|---|---|---|
| ACTA1 | Internal node | 0 | DEA |
| ACTA1_rna | Internal node | 0 | DEA |
| ACTG2 | Internal node | 0 | DEA |
| ACTG2_rna | Internal node | 0 | DEA |
| AKT1S1_phosphorylated | Internal node | 1 | DEA |
| AREG_Cytosol | Internal node | 0 | DEA |
| AREG_extracellular | Input node | 0 | DEA |
| AREG_rna | Internal node | 0 | DEA |
| AREG_Secreted_compartment | Internal node | 0 | DEA |
| ARHGEF7 | Internal node | 1 | DEA |
| BMP4 | Internal node | 1 | DEA |
| BMP4_rna | Internal node | 1 | DEA |
| CALML3 | Internal node | 0 | DEA |
| CASQ2 | Input node | 0 | DEA |
| CAV3 | Internal node | 0 | DEA |
| CAV3_rna | Internal node | 0 | DEA |
| CCL11 | Internal node | 1 | DEA |
| CCL11_rna | Internal node | 1 | DEA |
| CCL8 | Internal node | 0 | DEA |
| CCL8_rna | Internal node | 0 | DEA |
| CFL1 | Input node | 1 | 22954256 |
| Collagens_extracellular | Input node | 1 | 31521169 |
| CTGF_extracellular | Input node | 1 | 34108441 |
| CXCL1 | Internal node | 1 | DEA |
| CXCL1_rna | Internal node | 1 | DEA |
| CXCL12_extracellular | Input node | 1 | DEA |
| CXCL12_rna | Internal node | 1 | DEA |
| CXCL12_Secreted_compartment | Internal node | 1 | DEA |
| CXCL12CXCR4_complex | Internal node | 1 | DEA |
| CXCL2 | Internal node | 1 | DEA |
| CXCL2_rna | Internal node | 1 | DEA |
| EGF_extracellular | Input node | 1 | 35267539 |
| EGFR | Internal node | 1 | DEA |
| EGFR_complex | Internal node | 1 | DEA |
| EGFR_rna | Internal node | 1 | DEA |
| EMILIN1 | Internal node | 1 | DEA |
| FAP | Internal node | 1 | DEA |
| FAP_rna | Internal node | 1 | DEA |
| FGF1 | Input node | 1 | 33568624 |
| FGF2_extracellular | Input node | 1 | 32557854 |
| FGF3 | Input node | 1 | 35267539 |
| FGF4 | Input node | 1 | 33081025 |
| FGF7 | Internal node | 1 | DEA |
| FGF7_rna | Internal node | 1 | DEA |
| FGFRFGF1_complex | Internal node | 1 | DEA |
| FGFRFGF2_complex | Internal node | 1 | DEA |
| FGFRFGF3_complex | Internal node | 1 | DEA |
| FGFRFGF4_complex | Internal node | 1 | DEA |
| FN_extracellular | Input node | 1 | 35481621 |
| GAST | Input node | 1 | 28560291 |
| GASTCCKBR_complex | Internal node | 0 | DEA |
| GLI2 | Internal node | 1 | DEA |
| GLI2_rna | Internal node | 1 | DEA |
| GLNS | Input node | 1 | 27829138 |
| M_glc_D_e_simple_molecule | Input node | 1 | DEA |
| HGF_extracellular | Input node | 1 | 22282252 |
| HIF1A | Internal node | 1 | DEA |
| HRC | Input node | 0 | DEA |
| IFNG | Input node | 1 | 33005420 |
| IGF1_extracellular | Input node | 1 | DEA |
| IGF1_rna | Internal node | 1 | DEA |
| IGF1_Secreted_compartment | Internal node | 1 | DEA |
| IGF2_extracellular | Input node | 1 | DEA |
| IGF2_rna | Internal node | 1 | DEA |
| IGF2_Secreted_compartment | Internal node | 1 | DEA |
| IGFBP3 | Input node | 1 | DEA |
| IGFBP4 | Input node | 1 | 19536088 |
| IHH | Input node | 1 | 35224148 |
| IL12 | Input node | 0 | DEA |
| IL18 | Input node | 0 | 31231372 |
| IL1A | Input node | 1 | 31231372 |
| IL1B_extracellular | Input node | 1 | 31231372 |
| IL1R_complex_Cytosol | Internal node | 0 | DEA |
| IL1R_complex_Cytosol_2 | Internal node | 0 | DEA |
| IL6_extracellular | Input node | 1 | 35267539 |
| IRS2_phosphorylated | Internal node | 1 | DEA |
| ITGA11 | Internal node | 1 | DEA |
| ITGA11_rna | Internal node | 1 | DEA |
| ITGA11ITGB1_complex_Cytosol | Internal node | 1 | DEA |
| ITGA11ITGB1_complex_Cytosol_active | Internal node | 1 | DEA |
| ITGA1ITGB1_complex_Cytosol | Internal node | 0 | DEA |
| ITGA1ITGB1_complex_Cytosol_active | Internal node | 0 | DEA |
| ITGAVITGB6_complex_Cytosol | Internal node | 0 | DEA |
| ITGAVITGB6_complex_Cytosol_active | Internal node | 0 | DEA |
| Large_Latent_Complex_complex_extracellular | Input node | 1 | DEA |
| Large_Latent_Complex_complex_Secreted_compartment | Internal node | 1 | DEA |
| LGALS1 | Input node | 1 | 24229053 |
| LIF_extracellular | Input node | 1 | 34947829 |
| LIMK1_phosphorylated | Internal node | 1 | DEA |
| LOX | Internal node | 1 | DEA |
| LOX_rna | Internal node | 1 | DEA |
| LPA_simple_molecule | Input node | 0 | DEA |
| MAP3K7TAB_complex | Internal node | 1 | DEA |
| MIF | Input node | 0 | 24939415 |
| MIR101_antisense_rna | Input node | 0 | 28289080 |
| MIR141_antisense_rna | Input node | 0 | 28289080 |
| MIR155_antisense_rna | Input node | 1 | 23171795 |
| MIR200B_antisense_rna | Input node | 0 | 28289080 |
| MIR205_antisense_rna | Input node | 0 | 28289080 |
| MIR211_antisense_rna | Input node | 1 | 31702390 |
| MIR214_antisense_rna | Input node | 0 | 23171795 |
| MIR221_antisense_rna | Input node | 1 | 28289080 |
| MIR31_antisense_rna | Internal node | 1 | 28289080 |
| MMP13 | Internal node | 1 | DEA |
| MMP13_rna | Internal node | 1 | DEA |
| MMP14 | Internal node | 1 | DEA |
| MMP14_rna | Internal node | 1 | DEA |
| MMP2 | Internal node | 1 | DEA |
| MYLK_phosphorylated | Internal node | 0 | DEA |
| NDUFA4L2 | Internal node | 0 | DEA |
| NOX4 | Internal node | 1 | DEA |
| NOX4_rna | Internal node | 1 | DEA |
| OSM | Input node | 1 | 35192545 |
| PDGF_extracellular | Input node | 1 | 34272173 |
| PDGFPDGFRA_complex | Internal node | 1 | DEA |
| PDGFRA | Internal node | 1 | DEA |
| PDGFRA_rna | Internal node | 1 | DEA |
| PGE2_simple_molecule_extracellular | Input node | 1 | 33271839 |
| phospholipid_simple_molecule | Input node | 1 | |
| PI45P2_simple_molecule | Input node | 1 | 34108441 |
| PLAU | Internal node | 1 | DEA |
| PLAU_rna | Internal node | 1 | DEA |
| PLG | Input node | 1 | 33921488 |
| POSTN_extracellular | Input node | 1 | 35267539 |
| PPBP | Internal node | 0 | DEA |
| PPBP_rna | Internal node | 0 | DEA |
| proPLAU_extracellular | Input node | 1 | 24229053 |
| PTCH2_rna | Internal node | 1 | DEA |
| PTGS2 | Internal node | 1 | DEA |
| PTGS2_rna | Internal node | 1 | DEA |
| PTPN6 | Input node | 0 | DEA |
| RYR2TRDNASPH_complex | Input node | 0 | DEA |
| RYR2TRDNASPHHRCCASQ2_complex | Internal node | 0 | DEA |
| SEPTINE4 | Internal node | 0 | DEA |
| SERPINE1 | Internal node | 1 | DEA |
| SERPINE1_rna | Internal node | 1 | DEA |
| SHH | Input node | 1 | 28496132 |
| SMOPTCH_complex | Internal node | 1 | DEA |
| SMOX | Internal node | 1 | DEA |
| TGFB3_Cytosol | Internal node | 1 | DEA |
| TGFB3_extracellular | Internal node | 1 | DEA |
| TGFB3_rna | Internal node | 1 | DEA |
| TNF | Input node | 0 | DEA |
| VTN | Input node | 1 | 33211735 |
| WAS_phosphorylated | Internal node | 0 | DEA |
| WNT7 | Input node | 1 | 34108441 |
| WWTR1 | Input | 1 | DEA |
