## Supplementary material for "Hybrid computational modeling highlights reverse Warburg effect in breast cancer-associated fibroblasts": Supplementary_materials_TABLE-S2.pdf

**Table S2. Biological scenarios identified from breast cancer-specific peer-reviewed literature used for generic validation of the breast CAF-model's behavior in the Cell Collective platform.** Biological experimental conditions were used as the model's inputs and its outputs were compared to the biological outcome to confirm or refute a scenario's validation.

| Experimental observation | Reference | CAF model input | CAF model output | Coherence |
|---|---|---|---|---|
| A loss of Caveolin induces the metabolic reprogramming of breast CAFs by increasing mitochondrial dysfunction. | 22874531 | CAV-3 OFF/ON | CI, CIV OFF/ON | **YES** |
| Breast CAFs recombinantly expressing TGF-β show upregulation of myofibroblast markers β-actin. | | TGFB3 OFF/ON | Actin cytoskeletal OFF/ON | **YES** |
| Breast CAFs recombinantly expressing TGF-β show a loss of Cav-1/3 expression. | | TGFB3 OFF/ON | CAV3_rna ON/OFF | **YES** |
| Tumors derived from TGF-β ligand overexpressing breast CAFs display increased extracellular matrix deposition and increased secretion of: (a) Collagens, (b) Tenascin C. | | TGFB3 ON/OFF | COLLAGENS OFF/OFF | **NO** (Missing mechanistic information) |
| | | | TNC OFF/OFF | **NO** (Missing mechanistic information) |
| Breast CAFs promote tumor growth, in an angiogenesis-independent manner when overexpressing TGF-β3. | | TGFB3 OFF/ON | Tumor growth OFF/ON | **YES** |
| Upregulation of miR-221 in breast CAFs affects growth and migration by CTGF signaling pathway. | 29716623 | miR-221 ON | Tumor growth OFF/OFF | **NO** (Missing mechanistic information) |
| Downregulation of miR-320 in breast CAFs promotes tumor angiogenesis. | 22179046 | miR-320 OFF/ON | Angiogenesis OFF/ON | **YES** |
| miR-155 promotes proliferation of human breast CAFs. | 24152184 | miR-15 OFF/ON | Fibroblast proliferation OFF/ON | **YES** |
| Blockage of MAPK/p38 pathway diminished IL-32-induced tumor growth. | 30391782 | MAPK, p38 OFF/ON | Tumor growth OFF/ON | **YES** |
| Enhanced WNT expression in breast CAFs contributes to invasion and migration of breast cancer cells. | 32366558 | WNT7 OFF/ON | Migration into tumor OFF/ON | **YES** |
| In breast CAFs, IL-1β promotes cell invasion through IL-1R. | 27072893 | IL-1β OFF | Tumor invasion OFF/OFF | **NO** (Missing mechanistic information) |

| | | | | |
|---|---|---|---|---|
| CAFs' PDGF signaling has an active role in breast tumor progression. | 29380207 | PDGF OFF/ON | Tumor growth, migration OFF/ON | YES |
| In breast CAFs, CXCL12 promotes cell invasion through TGF-β pathway. | 31964880 | CXCL12 OFF/ON | Tumor invasion OFF/ON | YES |
| In breast CAFs, FGF2 promotes cancer growth and progression. | 32557854 | FGF2 OFF/ON | Tumor growth, growth factors OFF/ON | YES |
| CXCL1 is considered as a pro-inflammatory gene signature of breast CAFs. | 23831470 | CXCL1 OFF/ON | Cytokine production, immune system modulation OFF/ON | YES |
| Breast CAFs promote tumor growth through secretion of HGF. | 22282252 | HGF OFF/ON | Tumor growth OFF/ON | YES |
| miR-21 and miR-200B target TGF-β signaling and impact tumor progression and promotion in breast CAFs. | 27153853 | miR-21 OFF | | **NO** (Missing mechanistic information) |
| | | miR-200B OFF | | **NO** (Missing mechanistic information) |
| Myeloid-derived OSM reprograms breast CAFs to a more tumorigenic phenotype by eliciting the secretion of VEGF. | 35192545 | OSM OFF/ON | VEGF OFF/ON | YES |
| OSM promoted tumor growth through breast CAFs. | 35192545 | OSM OFF/ON | Tumor growth OFF/ON | YES |
| OSM induced the expression of classical CAF markers such as FAP, POSTN, VEGF, and IL6 in breast CAFs. | 35192545 | OSM OFF/ON | FAP, POSTN, VEGF, IL6 OFF/ON | YES |
| OSM induced signatures related to fibroblast activation and JAK/STAT3 signaling, in agreement with increased STAT3 phosphorylation by OSM in breast CAFs. | 35192545 | OSM OFF/ON | STAAT3 OFF/OFF | **NO** (Missing mechanistic information) |
| Plasmin expression affects tumor cell invasion. | 10190278 | Plasmin OFF/ON | Matrix degradation, matrix effects OFF/ON | YES |
| Plasmin expression is required for activation of EMT in breast cancer. | 19546228 | Plasmin ON/OFF | Matrix effect OFF/OFF | **NO** (Missing mechanistic information) |
| TGF-β promotes CXCL5 secretion in breast CAFs. | 32237072 | TGF-β ON/OFF | CXCL5 ON/OFF | YES |
| Lactate production in breast CAFs promotes breast cancer tumor growth. | 22129993 | Lactate OFF/ON | Tumor growth OFF/ON | YES |
| TFAM-deficient breast CAFs showed evidence of mitochondrial dysfunction. | 22129993 | TFAM OFF/ON | CI, CV OFF/ON | YES |
| SIRT3 in breast CAFs was found to:<br>(a) Suppress HIF-1α and its target genes; | 22589271; 22749020 | SIRT3 OFF/ON | HIF1A OFF/OFF | **NO** (Missing mechanistic |

| | | | | |
|---|---|---|---|---|
| (b) Suppress tumor growth and proliferation;<br>(c) Suppress ROS production. | | | | information) |
| | | | Tumor growth, tumor fibroblast proliferation OFF/ON | **YES** |
| | | | ROS production OFF/ON | **YES** |
| Breast CAFs promote tumor growth and angiogenesis through elevated CXCL12 secretion. | 15882617 | CXCL12 OFF/ON | Tumor growth, Angiogenesis OFF/ON | **YES** |
| LOX family members are considered as ECM-modifying enzymes in breast CAFs by remodeling the extracellular matrix. | 33037194 | LOX OFF/ON | ECM regulation phenotype and matrix effects OFF/ON | **YES** |
| Downregulation of MiR-205 in breast CAFs promotes VEGF-independent angiogenesis through activation of IL-11/IL-15 signaling by YAP1. | 29109792 | MiR-205 OFF | | **NO**<br>(Co-activator needed) |
| In breast cancer, HIF-1α transcriptionally upregulates glycolytic enzymes and lactate production. | 28623342 | HIF1A OFF/ON | Glycolytic enzymes, lactate OFF/ON | **YES** |
| Lactate generated by hypoxic breast CAFs promotes cell invasion. | 30799198 | Lactate OFF/ON | Tumor invasion OFF/ON | **YES** |
| Inhibiting NF-κB signaling in fibroblasts was shown to reduce inflammatory cytokine secretion. | 28378188 | NFkB OFF/ON | Cytokine production phenotype OFF/ON | **YES** |
| Ets2 inactivation through depletion of Pten in breast CAFs was sufficient to decrease tumor growth and progression. | 19847259 | PTEN OFF | | **NO**<br>(Co-activator needed) |
| SERPINE1 promotes cellular invasion in breast CAFs. | 33000256 | SERPINE1 OFF/ON | Tumor invasion OFF/ON | **YES** |
| CASQ2 overexpression accelerated tumorigenesis, induced collagen structure remodeling, and increased distant metastasis. | 34743414 | CASQ2 OFF/ON | Collagens OFF/OFF | **NO** (Missing mechanistic information) |
| Hypoxic breast CAFs led to sustained elevation of HIF-1α. | 32492417 | Hypoxia OFF/ON | HIF1A OFF/ON | **YES** |
