## Supplementary material for "Hybrid computational modeling highlights reverse Warburg effect in breast cancer-associated fibroblasts": Supplementary_materials_TABLE-S4.pdf

**Table S4. Metabolic enzymes with projected maximal regulatory trap-space equal to 0 in breast CAF-specific initial conditions and their associated catalyzed reaction constrained to 0 in MitoCore.**

| Enzyme | Complete Name | Metabolic subsystem | Catalyzed reaction | Detailed reaction |
|---|---|---|---|---|
| GLUNm | Mitochondrial Glutaminase | Glutamine degradation | R_GLUNm | L-Glutamine[m] + $H_2O$[m] + $NAD^+$[m] → L-Glutamate[m] + $NH_4$[m] |
| CI_MitoCore | NADH Dehydrogenase | Electron transport chain | R_CI_MitoCore | $H^+$[m] + NADH[m] + Ubiquinone[m] ⇌ $NAD^+$[m] + $H^+$[m] + Ubiquinol[m] |
| HMGCOASim | Hydroxymethylglutaryl Coenzyme A Synthase | Ketogenesis | R_ HMGCOASim | Acetyl-CoaA[m] + Acetoacetyl-CoA[m] + $H_2O$[m] ⇌ 3-Hydroxy-3-methylglutaryl-CoA[m] + CoA[m] |
| CIII_MitoCore | Cytochrome C Reductase | Electron transport chain | R_CIII_MitoCore | $H^+$[m] + Ubiquinol[m] + Ferricytochrome C[m] ⇌ $H^+$[m] + Ubiquinone[m] + Ferrocytochrome C[m] |
| CIV_MitoCore | Cytochrome C Oxidase | Electron transport chain | R_CIV_MitoCore | $H^+$[m] + $O_2$[m] + Ferrocytochrome C[m] → $H^+$[m] + $H_2O$[m] + Ferricytochrome C[m] |
| ICDHxm | Isocitrate Dehydrogenase | Tricarboxylic acid cycle | R_ICDHxm | Isocitrate[m] + $NAD^+$[m] → 2-Oxoglutarate[m] + $CO_2$[m] + NADH[m] |
| PDHm | Pyruvate Dehydrogenase | Tricarboxylic acid cycle | R_PDHm | Pyruvate[m] + CoA[m] + $NAD^+$[m] → Acetyl-CoA[m] + $CO_2$[m] + NADH[m] + $H^+$[m] |
