## Supplementary material for "Hybrid computational modeling highlights reverse Warburg effect in breast cancer-associated fibroblasts": Supplementary_materials_TABLE-S5.pdf

**Table S5. Metabolic enzymes with projected maximal regulatory trap-space equal to 0 in breast CAF-specific initial conditions and their associated catalyzed reaction constrained to 0 in MitoCore.**

| Metabolite | Complete name | Metabolic subsystem | Producing reaction | Detailed reaction |
|---|---|---|---|---|
| M_gln_L_c | L-Glutamine [cytoplasmic] | Glutamine synthesis | R_GLNS | ATP [c] + L-Glutamate[c] + $NH_4$[c] → **L-Glutamine [c]** + ADP[c] + Orthophosphate[c] |
| | | L-Glutamine transport | R_r2525 | L-Glutamine[e] ⇌ **L-Glutamine [c]** |
| M_hmgcoa_m | 3-Hydroxy-3-methylglutaryl-CoA [mitochondrial] | Leucine degradation | R_MGCHrm | 3-Methylglutaconyl-CoA[m] + $H_2O$[m] ⇌ **3-Hydroxy-3-methylglutaryl-CoA**[m] |
| | | Ketogenesis | R_ HMGCOASim | Acetyl-CoaA[m] + Acetoacetyl-CoA[m] + $H_2O$[m] ⇌ **3-Hydroxy-3-methylglutaryl-CoA**[m] + CoA[m] |
| M_accoa_m | Acetyl-CoA [mitochondrial] | Isoleucine degradation | R_ACACT10m | CoA[m] + 2-Methylacetoacetyl-CoA[m] → Propanoyl-CoA[m] + **Acetyl-CoA[m]** |
| | | FA and ketone body metabolism, ketogenesis | R_ACACT1rm | **Acetyl-CoA[m]** ⇌ CoA[m] + Acetoacetyl-CoA[m] |
| | | TCA cycle periphery | R_ACITLm_MitoCore | ATP[m] + Citrate[m] + CoA[m] → ADP[m] + Orthophosphate[m] + **Acetyl-CoA[m]** + Oxaloacetate[m] |
| | | Alcohol metabolism | R_ACSm | ATP[m] + Acetate[m] + CoA[m] → AMP[m] + Diphosphate[m] + **Acetyl-CoA[m]** |
| | | Ketogenesis / Leucine degradation | R_HMGLm | 3-Hydroxy-3-methylglutaryl-CoA[m] → **Acetyl-CoA[m]** + Acetoacetate[m] |
| | | Beta-alanine degradation | R_MMSAD3m | 3-Oxopropanoate[m] + CoA[m] + $NAD^+$[m] → **Acetyl-CoA[m]** + $CO_2$[m] + NADH[m] + $H^+$[m] |
| | | Beta-alanine degradation | R_MMSAD3m2_MitoCore | 3-Oxopropanoate[m] + CoA[m] + $NADP^+$[m] → **Acetyl-CoA[m]** + $CO_2$[m] + NADPH[m] + $H^+$[m] |
| | | Fatty acid metabolism | R_MTPC14_MitoCore | Trans-Hexadec-2-enoyl-CoA[m] + $H_2O$[m] + $NAD^+$[m] + CoA[m] ⇌ Lauroyl-CoA[m] + NADH[m] + **Acetyl-CoA[m]** |

| | | | | |
|---|---|---|---|---|
| | | | R_MTPC16_MitoCore | Trans-Hexadec-2-enoyl-CoA[m] + $H_2O$[m] + $NAD^+$[m] + CoA[m] ⇌ Tetradecanoyl-CoA[m] + NADH[m] + **Acetyl-CoA[m]** |
| | | | R_r0287 | CoA[m] + 3-Oxohexanoyl-CoA[m] ⇌ **Acetyl-CoA[m]** + Butanoyl-CoA[m] |
| | | | R_r0634 | Octanoyl-CoA[m] + **Acetyl-CoA[m]** ⇌ CoA[m] + 3-Oxodecanoyl-CoA[m] |
| | | | R_r0724 | Decanoyl-CoA[m] + **Acetyl-CoA[m]** ⇌ CoA[m] + 3-Oxododecanoyl-CoA[m] |
| | | | R_r0732 | CoA[m] + 3-Oxooctanoyl-CoA[m] ⇌ Hexanoyl-CoA[m] + **Acetyl-CoA[m]** |
| | | Glycolysis, gluconeogenesis | R_PDHm | Pyruvate[c] + CoA[c] + $NAD^+$[c] → **Acetyl-CoA[c]** + $CO_2$[c] + NADH[c] + $H^+$[c] |
| M_akg_m | 2-Oxoglutarate [mitochondrial] | Lysine degradation | R_2AMADPTmC_MitoCore | **2-Oxoglutarate[m]** + L-Aminoadipate[m] → 2-Oxoglutarate[c] + L-Aminoadipate[c] |
| | | Tryptophan Metabolism | R_2OXOADPTmC_MitoCore | **2-Oxoglutarate[m]** + 2-Oxoadipate[c] → 2-Oxoglutarate[c] + 2-Oxoadipate[m] |
| | | Malate aspartate shuttle | R_ASPTAm | L-Aspartate[m] + **2-Oxoglutarate[m]** ⇌ Oxaloacetate[m] + L-Glutamate[m] |
| | | Glutamate degradation and synthesis | R_GLUDxm | L-Glutamate[m] + $NAD^+$[m] + $H_2O$[m] → **2-Oxoglutarate[m]** + $NH_4$[m] + NADH[m] + $H^+$[m] |
| | | | R_GLUDym | L-Glutamate[m] + $NADP^+$[m] + $H_2O$[m] → **2-Oxoglutarate[m]** + $NH_4$[m] + NADPH[m] + $H^+$[m] |
| | | Tricarboxylic acid cycle | R_ICDHxm | Isocitrate[m] + $NAD^+$[m] → **2-Oxoglutarate[m]** + $CO_2$[m] + NADH[m] |
| | | | R_ICDHyrm | Isocitrate[m] + $NADP^+$[m] ⇌ **2-Oxoglutarate[m]** + $CO_2$[m] + NADPH[m] |
| M_cit_m | Citrate [mitochondrial] | Mitochondrial transporters | R_CITtamB | Citrate[c] + L-Malate[m] → **Citrate[m]** + L-Malate[c] |
| | | | R_CITtbm | Citrate[c] + Phosphoenolpyruvate[m] → **Citrate[m]** + Phosphoenolpyruvate[c] |
| | | | R_r0917 | Citrate[c] + Isocitrate[m] → **Citrate[m]** + Isocitrate[c] |
| | | | R_r0917b_MitoCore | Citrate[c] + Isocitrate[m] → **Citrate[m]** + Isocitrate[c] |

| | | | | |
|---|---|---|---|---|
| | | Tricarboxylic acid cycle | R_CSm | Acetyl-CoA[m] + $H_2O$[m] + Oxaloacetate[m] → **Citrate[m]** + CoA[m] |
| M_fum_m | Fumarate [mitochondrial] | Electron transport chain | R_CII_MitoCore | FAD[m] + Succinate[m] ⇌ **Fumarate[m]** + $FADH_2$[m] |
| | | Mitochondrial transporters | R_FUMtmB_Mitocore | Fumarate[c] + Pi[m] → **Fumarate[m]** + Pi[c] |
| M_icit_m | Isocitrate [mitochondrial] | Tricarboxylic acid cycle | R_ACONTm | Citrate[m] ⇌ **Isocitrate[m]** |
| M_mal_L_m | L-Malate [mitochondrial] | Malate aspartate shuttle | R_AKGMALtm | L-Malate[c] + 2-Oxoglutarate[m] ⇌ **L-Malate[m]** + 2-Oxoglutarate[c] |
| | | Mitochondrial transporters | R_MALSO3tm | L-Malate[c] + Sulfite[m] ⇌ **L-Malate[m]** + Sulfite[c] |
| | | | R_MALSO4tm | L-Malate[c] + Sulfate[m] ⇌ **L-Malate[m]** + Sulfate[c] |
| | | | R_MALTSULtm | L-Malate[c] + Thiosulfate[m] ⇌ **L-Malate[m]** + Thiosulfate[c] |
| | | | R_r0913 | L-Malate[c] + Isocitrate[m] → **L-Malate[m]** + Isocitrate[c] |
| | | Tricarboxylic acid cycle | R_FUMm | Fumarate[m] + $H_2O$[m] ⇌ **L-Malate[m]** |
| M_oaa_m | Oxaloacetate [mitochondrial] | Tricarboxylic acid cycle periphery | R_ACITLm_MitoCore | ATP[m] + Citrate[m] + CoA[m] → ADP[m] + Orthophosphate[m] + Acetyl-CoA[m] + **Oxaloacetate[m]** |
| | | | R_MDHm | L-Malate[m] + $NAD^+$[m] ⇌ **Oxaloacetate[m]** + NADH[m] + $H^+$[m] |
| | | | R_PCm | ATP[m] + Pyruvate[m] + $HCO3^-$[m] → ADP[m] + Orthophosphate[m] + **Oxaloacetate[m]** |
| M_gln_L_m | L-Glutamine [mitochondrial] | Mitochondrial transporters | R_GLNtm | L-Glutamine[c] ⇌ **L-Glutamine**[m] |
| M_succ_m | Succinate [mitochondrial] | Ketone bodies degradation | R_OCOAT1m | Succinyl-CoA[m] + Acetoacetate[m] ⇌ **Succinate[m]** + Acetoacetyl-CoA[m] |
| | | Mitochondrial transporters | R_SUCCt2m | Succinate[c] + Pi[m] ⇌ **Succinate[m]** + Pi[c] |
| | | | R_SUCCt3m_MitoCore | Succinate[c] + L-Malate[m] ⇌ **Succinate[m]** + L-Malate[c] |

| | | | | |
|---|---|---|---|---|
| | | | R_r0829 | Succinate[c] + Sulfate[m] ⇌ **Succinate[m]** + Sulfate[c] |
| | | | R_r0830 | Succinate[c] + Sulfite[m] ⇌ **Succinate[m]** + Sulfite[c] |
| | | | R_r0830B_MitoCore | Succinate[c] + Thiosulfate[m] ⇌ **Succinate[m]** + Thiosulfate[c] |
| | | Tricarboxylic acid cycle | R_SUCOAS1m | GDP[m] + Orthophosphate[m] + Succinyl-CoA[m] ⇌ GTP[m] + **Succinate[m]** + CoA[m] |
| | | | R_SUCOASm | GDP[m] + Succinyl-CoA[m] ⇌ GTP[m] + **Succinate[m]** |
| | | GABA shunt | R_r0178 | Succinate semialdehyde[m] + $NAD^+$[m] + $H_2O$[m] → **Succinate[m]** + NADH[m] + $H^+$[m] |
| M_succoa_m | Succinyl-CoA [mitochondrial] | Tricarboxylic acid cycle | R_AKGDm | 2-Oxoglutarate[m] + CoA[m] + $NAD^+$[m] → **Succinyl-CoA[m]** + $CO_2$[m] + NADH[m] + $H^+$[m] |
| | | Propanoate metabolism | R_MMMm | (R)-Methylmalonyl-CoA[m] ⇌ **Succinyl-CoA[m]** |
| M_acac_m | Acetoacetic acid [mitochondrial] | Mitochondrial transporters | R_ACACt2mB_MitoCore | Acetoacetic acid[c] + $H^+$[c] ⇌ **Acetoacetic acid[m]** + $H^+$[m] |
| | | Ketone bodies degradation | R_BDHm | 3-Hydroxybutyric acid[m] + $NAD^+$[m] ⇌ **Acetoacetic acid[m]** + $H^+$[m] + NADH[m] |
| | | Ketogenesis, leucine degradation | R_HMGLm | 3-Hydroxy-3-methylglutaryl-CoA[m] → **Acetoacetic acid[m]** + Acetyl-CoA[m] |
| M_glu_L_m | L-Glutamate [mitochondrial] | GABA shunt | R_ABTArm | 4-aminobutanoate[m] + 2-oxoglutarate[m] ⇌ **L-Glutamate[m]**+ Succinate Semialdehyde[m] |
| | | Beta-alanine degradation | R_APAT2rm | 2-oxoglutarate[m] + Beta-alanine[m] ⇌ **L-Glutamate[m]** + 3-Oxopropanoate[m] |
| | | Malate asparte shuttle | ASPGLUmB_MitoCore | L-aspartate[m] + L-glutamate[c] + $H^+$[c] → L-aspartate[c] + **L-Glutamate[m]** + $H^+$[m] |
| | | Malate aspartate shuttle | R_ASPTAm | L-Aspartate[m] + 2-Oxoglutarate[m] ⇌ Oxaloacetate[m] + **L-Glutamate[m]** |
| | | Cysteine degradation | R_CYSTAm | 2-Oxoglutarate[m] + L-Cysteine[m] ⇌ **L-Glutamate[m]** + Mercaptopyruvate[m] |

| | | | | |
|---|---|---|---|---|
| | | Mitochondrial transporters | R_GLUt2mB_MitoCore | L-Glutamate[c] + $H^+$[c]] ⇌ **L-Glutamate[m]** + $H^+$[m] |
| | | Isoleucine degradation | R_ILETAm | 2-Oxoglutarate[m] + L-isoleucine[m] ⇌ **L-Glutamate[m]** + 3-Methyl-2-oxopentanoicacid[m] |
| | | Leucine degradation | R_LEUTAm | 2-Oxoglutarate[m] + L-Leucine[m] ⇌ **L-Glutamate[m]** + 4-Methyl-2-oxopentanoate[m] |
| | | Ornithine degradation | R_ORNTArm | 2-Oxoglutarate[m] + L-Ornithine[m] ⇌ **L-Glutamate[m]** + L-Glutamate 5-semialdehyde[m] |
| | | Valine degradation | R_VALTAm | 2-Oxoglutarate[m] + L-Valine[m] ⇌ **L-Glutamate[m]** + 3-Methyl-2-oxobutanoicacid[m] |
| | | Proline, ornithine degradation | R_r0074 | L-Glutamate 5-semialdehyde[m] + H2O[m] + NAD+[m] ⇌ **L-Glutamate[m]** + 2 $H^+$[m] + NADH[m] |
| | | Tricarboxylic acid cycle periphery | R_r0081 | 2-Oxoglutarate[m] + L-Alanine[m] ⇌ **L-Glutamate[m]**+ Pyruvate[m] |
| | | Lysine degradation | R_r0450 | 2-Oxoglutarate[m] + L-2-Aminoadipate[m] ⇌ **L-Glutamate[m]** + 2-Oxodipate[m] |
| | | Lysine degradation | R_r0525 | H2O[m] + NAD+[m] + Saccharopine[m] ⇌ **L-Glutamate[m]** + $H^+$[m] + Allysine[m] + NADH[m] |
| M_bhb_m | 3-Hydroxybutyric acid [mitochondrial] | Mitochondrial transporters | R_BHBtmB_MitoCore | 3-Hydroxybutyric acid[c] + $H^+$[c] ⇌ **3-Hydroxybutyric acid[m]** + $H^+$[m] |
