## Supplementary material for "Hybrid computational modeling highlights reverse Warburg effect in breast cancer-associated fibroblasts": Supplementary_materials_TABLE-S7.pdf

**Table S7. Metabolic flux distribution in breast CAF conditions with maximal ATP production as objective function.**

| Reaction | Flux |
|---|---|
| EX_2hb_e | 0,01 |
| EX_ac_e | 0,01 |
| EX_acac_e | 0 |
| EX_akg_e | 0 |
| EX_ala_B_e | 0 |
| EX_ala_L_e | -0,01 |
| EX_arg_L_e | 0 |
| EX_argsuc_e | 0 |
| EX_asn_L_e | -0,01 |
| EX_asp_L_e | -0,154 |
| EX_bhb_e | 0 |
| EX_bilirub_e | 0 |
| EX_biomass_e | 0 |
| EX_but_e | -7,98E-17 |
| EX_chol_e | -2,15E-16 |
| EX_cit_e | 0 |
| EX_citr_L_e | 0 |
| EX_co_e | 0 |
| EX_co2_e | 0,41 |
| EX_creat_e | 0 |
| EX_cyan_e | 0 |
| EX_cys_L_e | -0,001 |
| EX_etoh_e | 0 |
| EX_fe2_e | 0 |
| EX_for_e | 0 |
| EX_fum_e | 0 |
| EX_glc_D_e | -0,9 |
| EX_gln_L_e | 0 |
| EX_glu_L_e | 0 |
| EX_gly_e | -3,42E-17 |
| EX_glyc_e | -0,01 |
| EX_h_e | 1,402 |
| EX_h2o_e | 0 |
| EX_HC00250_e | 0 |
| EX_hco3_e | -0,236 |
| EX_hdca_e | 1,61E-17 |
| EX_his_L_e | -0,01 |
| EX_icit_e | 0 |
| EX_ile_L_e | 0 |
| EX_lac_L_e | 1,995 |
| EX_leu_L_e | 0 |
| EX_lys_L_e | 6,16E-33 |
| EX_mal_L_e | 0 |
| EX_mercplac_e | 0 |
| EX_met_L_e | 0 |
| EX_nad_e | 0 |
| EX_nadh_e | 0 |
| EX_nh4_e | 0,225 |
| EX_no_e | 0 |

| | |
|---|---|
| EX_o2_e | -0,001 |
| EX_oaa_e | 1,28E-17 |
| EX_pchol_hs_e | 1,47E-16 |
| EX_pcreat_e | 0 |
| EX_pe_hs_e | 9,37E-17 |
| EX_phe_L_e | 0 |
| EX_pi_e | 1,91E-17 |
| EX_ppa_e | 0 |
| EX_pro_L_e | -1,25E-18 |
| EX_ps_hs_e | 7,98E-17 |
| EX_ser_L_e | 4,07E-17 |
| EX_so3_e | 0,001 |
| EX_succ_e | 0 |
| EX_tcynt_e | 0 |
| EX_thr_L_e | -0,01 |
| EX_trp_L_e | 0 |
| EX_tsul_e | 0 |
| EX_tyr_L_e | 8,13E-20 |
| EX_urea_e | 0 |
| EX_val_L_e | 0 |
| EX_fol_e | -1,54E-17 |
| OF_ATP_MitoCore | 0 |
| OF_HEME_MitoCore | 0 |
| OF_LIPID_MitoCore | 0 |
| OF_PROTEIN_MitoCore | 0 |
| HEX1 | 0,9 |
| G6PPer | 0 |
| PGI | 0,9 |
| PFK | 0,9 |
| FBP | 0 |
| FBA | 0,9 |
| TPI | 0,91 |
| GAPD | 1,81 |
| PGK | 1,81 |
| PGM | 1,81 |
| ENO | 1,81 |
| PYK | 1,984 |
| r0122 | 0 |
| PEPCK | 0,174 |
| LDH_L | 1,995 |
| G6PDH2r | 0 |
| PGL | 0 |
| GND | 0 |
| RPI | 0 |
| RPE | 0 |
| TKT1 | 0 |
| TALA | 0 |
| TKT2 | 0 |
| PDHm | 0 |
| CSm | 0 |
| ACONTm | 0 |
| ICDHxm | 0 |
| ICDHyrm | 0 |

| | |
|---|---|
| AKGDm | 0 |
| SUCOAS1m | 0 |
| SUCOASm | 0 |
| FUMm | 0 |
| MDHm | 0 |
| CI_MitoCore | 0 |
| CII_MitoCore | 0 |
| CIII_MitoCore | 0 |
| CIV_MitoCore | 0 |
| CV_MitoCore | 0,666666667 |
| PEPCKm | 0 |
| PCm | 0 |
| ME2m | 0 |
| ME1m | 0 |
| r0081 | 0 |
| ACITLm_MitoCore | 0 |
| NDPK1m | 0 |
| NNT_MitoCore | 0 |
| ADK1m | 1,115333333 |
| ME2 | 0 |
| ALATA_L | 0,01 |
| NDPK1 | 0,174 |
| FUM | 0 |
| ADK1 | 8,55E-17 |
| ICDHy | -0,01 |
| ACONT | -0,01 |
| ACITL | 0,01 |
| ASPTA | 0,164 |
| MDH | 0 |
| AKGMALtm | 0 |
| ASPGLUmB_MitoCore | 0 |
| ASPTAm | 0 |
| G3PD1 | 0,01 |
| r0205 | 0 |
| FACOAL160i | 8,55E-17 |
| C160CPT1 | 8,55E-17 |
| PPA | 8,55E-17 |
| r2435 | 8,55E-17 |
| C160CPT2 | 8,55E-17 |
| PPAm | 1,115333333 |
| ACOT2_MitoCore | 0 |
| ACADLC16_MitoCore | 0 |
| MECR16C_MitoCore | -1,54E-17 |
| MTPC16_MitoCore | 1,54E-17 |
| ACADLC14_MitoCore | 0 |
| MECR14C_MitoCore | -1,54E-17 |
| MTPC14_MitoCore | 1,54E-17 |
| r1447 | 1,54E-17 |
| r0638 | 0 |
| r0660 | -1,54E-17 |
| r0722 | 1,54E-17 |
| r0724 | -1,54E-17 |
| r1451 | 1,54E-17 |

| | |
|---|---|
| r0735 | 0 |
| r0728 | 7,96E-18 |
| r0726 | -7,96E-18 |
| r0634 | 7,96E-18 |
| r1448 | -7,96E-18 |
| r0633 | 0 |
| r0731 | 4,15E-17 |
| r0730 | 4,15E-17 |
| r0732 | 4,15E-17 |
| r1450 | 4,15E-17 |
| r0791 | 0 |
| r0734 | 4,15E-17 |
| r0733 | 4,15E-17 |
| r0287 | -7,98E-17 |
| r1446 | 0 |
| ECOAH1m | 0 |
| HACD1m | 0 |
| ACACT1rm | 0 |
| ACCOAC | 0 |
| MCOATA | 0 |
| ACOATA | 0 |
| r0678 | 0 |
| r0691 | 0 |
| r0681 | 0 |
| r0682 | 0 |
| r0760 | 0 |
| r0761 | 0 |
| r0762 | 0 |
| r0763 | 0 |
| r0764 | 0 |
| r0694 | 0 |
| r0695 | 0 |
| r0765 | 0 |
| r0766 | 0 |
| r0692 | 0 |
| r0693 | 0 |
| r0767 | 0 |
| r0768 | 0 |
| r0769 | 0 |
| r0770 | 0 |
| r0712 | 0 |
| r0713 | 0 |
| r0701 | 0 |
| r0702 | 0 |
| r0771 | 0 |
| r0772 | 0 |
| r0696 | 0 |
| r0697 | 0 |
| r0773 | 0 |
| FA160ACPH | 0 |
| FACOAL40im | 7,98E-17 |
| BDHm | 0 |
| OCOAT1m | 0 |

| | |
|---|---|
| HMGCOASim | 0 |
| HMGLm | 0 |
| LEUTAm | 0 |
| OIVD1m | 0 |
| r0655 | 0 |
| MCCCrm | 0 |
| MGCHrm | 0 |
| ILETAm | 0 |
| OIVD3m | 0 |
| r0603 | 0 |
| ECOAH9m | 0 |
| HACD9m | 0 |
| ACACT10m | 0 |
| VALTAm | 0 |
| OIVD2m | 0 |
| r0560 | 0 |
| ECOAH12m | 0 |
| 3HBCOAHLm | 0 |
| HIBDm | 0 |
| ACCOALm | 1,115333333 |
| MMSAD1m | 0 |
| PPCOACm | 0,236 |
| MMEm | 0 |
| MMMm | 0 |
| MMCDm | 0,236 |
| RE2649M | 1,115333333 |
| THRD_L | 0,01 |
| r1155 | 0 |
| r1154 | 0 |
| 2HBO | -0,01 |
| METAT | 0 |
| METAT2_MitoCore | 0 |
| AHC | 0 |
| ADNK1 | 0 |
| CYSTS | 0 |
| CYSTGL | 0 |
| CYSO | 0,001 |
| 3SALATAi | 0,001 |
| 3SPYRSP | 0,001 |
| CYSTA | 0 |
| CYSTAm | 0 |
| MCPST | 0 |
| MCPSTm_MitoCore | 0 |
| r0595m_MitoCore | 0 |
| r0595B_MitoCore | 0 |
| MCLOR | 0 |
| r0193 | 0 |
| TRPO2 | 0 |
| FKYNH | 0 |
| KYN3OX | 0 |
| HKYNH | 0 |
| 3HAO | 0 |
| PCLAD | 0 |

| | |
|---|---|
| r0645 | 0 |
| AMCOXO | 0 |
| AMCOXO2_MitoCore | 0 |
| 2OXOADPTmB_MitoCore | 0 |
| 2OXOADPTmC_MitoCore | 0 |
| 2OXOADOXm | 0 |
| r0541 | 0 |
| SACCD3m | 0 |
| r0525 | 0 |
| AASAD3m | 0 |
| R03103_MitoCore | 0 |
| r0450 | 0 |
| LYSOXc_MitoCore | -6,16E-33 |
| PPD2CSPc_MitoCore | 0 |
| 1PPDCRc_MitoCore | 0 |
| 1PPDCRc_NADPH_MitoCore | 0 |
| LPCOXc_MitoCore | 0 |
| RE1254C | 0 |
| r0594 | 0 |
| 2AMADPTmB_MitoCore | 0 |
| 2AMADPTmC_MitoCore | 0 |
| PROD2mB_MitoCore | -1,54E-17 |
| G5SADrm | 0 |
| r0074 | 0 |
| GLU5Km | 0 |
| G5SDym | 0 |
| P5CRm | 0 |
| P5CRxm | -1,67E-17 |
| ORNTArm | 0 |
| ORNDC | 0 |
| PTRCOX1 | 0 |
| r0464c_MitoCore | 0 |
| ABUTD | 0 |
| ARGDCm | 0 |
| AGMTm | 0 |
| PTRCAT1m_MitoCore | 0 |
| APRTO2m_MitoCore | 0 |
| NABTNOm | 0 |
| 4aabutn_MitoCore | 0 |
| GLUDC | 0 |
| 4ABUTtm | 0 |
| ABTArm | 0 |
| r0178 | 0 |
| GLUDxm | 0 |
| GLUDym | 0 |
| GLUDxi | 0,185 |
| GLUDy | 0 |
| GLNS | 0 |
| GLUNm | 0 |
| GLUN_MitoCore | 0 |
| PGCD | 0 |
| PSERT | 0 |
| PSP_L | 0 |

| | |
|---|---|
| GHMT2r | -6,94E-18 |
| FOLR2 | 0 |
| DHFR | 0 |
| MTHFD | -6,94E-18 |
| MTHFC | 0,01 |
| FTCD | 0,01 |
| FTHFL | 0 |
| FTHFDH | 0,01 |
| r0060 | 0 |
| GHMT2rm | -4,12E-17 |
| GCCam | 1,84E-17 |
| GCCbim | 1,60E-17 |
| GCCcm | 1,60E-17 |
| r0514 | 1,54E-17 |
| r0226 | 1,54E-17 |
| MTHFDm | 0 |
| MTHFD2m | -5,85E-17 |
| MTHFCm | 3,47E-18 |
| FTHFLm | 0 |
| FTHFDHm_MitoCore | 0 |
| GLYATm | 0 |
| AOBUTDsm | 0 |
| AACTOORm_MitoCore | 0 |
| LGTHLm_MitoCore | 0 |
| GLYOXm | 0 |
| LDH_Dm_MitoCore | 0 |
| CBPSam | 0 |
| OCBTm | 0 |
| NOS1 | 0 |
| NOS2 | 0 |
| r0129 | 0 |
| AMPTASECG | 0 |
| GLUCYS | 0 |
| GTHS | 0 |
| r0399 | 0 |
| DHPR | 0 |
| TYRTA | 0 |
| TYRTB_MitoCore | 0 |
| 34HPPOR | 0 |
| HGNTOR | 0 |
| MACACI | -2,71E-20 |
| FUMAC | -2,71E-20 |
| ASNS1 | 0 |
| r0127 | 0,01 |
| HISD | 0,01 |
| URCN | 0,01 |
| IZPN | 0,01 |
| GluForTx | 0,01 |
| APAT2rm | 0 |
| MMSAD3m | 0 |
| MMSAD3m2_MitoCore | 0 |
| ASP1DC | 0 |
| CKc | 0 |

| | |
|---|---|
| CK | 0 |
| ACOAHi | 0,01 |
| ALCD2yf | 0 |
| ALCD2if | 0 |
| ACALDtm | 0 |
| ALDD2xm | 0 |
| ALDD2x | 0 |
| ACSm | 0 |
| ACS | 0 |
| ADSL1 | 0 |
| ADSS | 0 |
| AMPD1 | 0 |
| ARGN | 0 |
| ARGSL | 0 |
| ARGSS | 0 |
| ARGNm | 0 |
| ALASm | 0 |
| 5AOPtm | 0 |
| PPBNGS | 0 |
| HMBS | 0 |
| UPP3S | 0 |
| UPPDC1 | 0 |
| CPPPGO | 0 |
| PPPGOmB_MitoCore | 0 |
| FCLTm | 0 |
| PHEMEtm | 0 |
| HOXG | 0 |
| BILIRED | 0 |
| BILIRED2_MitoCore | 0 |
| PCHOLPm_hs | -2,15E-16 |
| GLYK | 0,01 |
| GLYC3Ptm | -9,71E-17 |
| GPAMm_hsB_MitoCore | 0 |
| AGPAT1B_MitoCore | 0 |
| CDSm | -9,71E-17 |
| PGPPTm | -9,71E-17 |
| PGPP_hsm_MitoCore | -9,71E-17 |
| CLS_hsm_MitoCore | 0 |
| CLPN_MitoCore | -4,86E-17 |
| CYTK1m | -9,71E-17 |
| NDPK3m | -9,71E-17 |
| SPODMm | 0 |
| GTHP | 0 |
| GTHPm | 0 |
| GTHO | -6,16E-33 |
| GTHOm | 0 |
| CITtamB | 0 |
| r0913 | 0 |
| CITtbm | 0 |
| r0917 | 0 |
| r0917b_MitoCore | 0 |
| PIt2mB_MitoCore | -1,8 |
| ATPtmB_MitoCore | -1,8 |

| | |
|---|---|
| HtmB_MitoCore | 0 |
| MALtm | 0 |
| MALSO3tm | 0 |
| MALTSULtm | 0 |
| MALSO4tm | 0 |
| SUCCt2m | 0 |
| r0830 | 0 |
| r0830B_MitoCore | 0 |
| r0829 | 0 |
| SUCCt3m_MitoCore | 0 |
| COAtmB_MitoCore | 0 |
| COAtmC_MitoCore | 0 |
| GLUt2mB_MitoCore | 0 |
| ORNt4mB_MitoCore | 0 |
| r2398B_MitoCore | 0 |
| r2402B_MitoCore | 0 |
| LYStmB_MitoCore | 0 |
| ORNt3mB_MitoCore | 0 |
| ARGtmB_MitoCore | 0 |
| r1427 | 0 |
| PYRt2m | 0 |
| ACACt2mB_MitoCore | 0 |
| FE2tm | 0 |
| ASNtm | 0 |
| r1437 | 0 |
| GLNtm | 0 |
| PROtm | 1,25E-18 |
| r1078 | 0 |
| r1436 | 0 |
| r1455 | 0 |
| TRPtm_MitoCore | 0 |
| GLYtm | 5,00E-17 |
| ILEt5m | 0 |
| LEUt5m | 0 |
| VALt5m | 0 |
| r1434 | 0 |
| r1435 | -3,37E-17 |
| r1440 | 0 |
| BALAtmr | 0 |
| UREAtm | 0 |
| FUMtmB_MitoCore | 0 |
| BHBtmB_MitoCore | 0 |
| PPAtmB_MitoCore | 0 |
| BUTt2mB_MitoCore | 7,98E-17 |
| FORt2mB_MitoCore | 0 |
| r0962B_MitoCore | 1,54E-17 |
| CHLtmB_MitoCore | 2,15E-16 |
| CO2tm | -0,236 |
| H2Otm | 1,564 |
| O2tm | 0 |
| GLYCtm | 4,86E-17 |
| CYANtm | 0 |
| TCYNTtmB_MitoCore | 0 |

| | |
|---|---|
| CREATtmdiffir | 0 |
| PCREATtmdiffirB_MitoCore | 0 |
| r0941 | 0,236 |
| r0838B_MitoCore | -1,84E-17 |
| Biomasst_MitoCore | 0 |
| PCFLOPm | -1,47E-16 |
| PSFLIPm | -7,98E-17 |
| PEFLIPm | -9,37E-17 |
| Biomass_MitoCore | 0 |
| O2t | 0,001 |
| CO2t | -0,41 |
| HCO3t_MitoCore | 0,236 |
| GLCt1r | 0,9 |
| HDCAtr | -1,61E-17 |
| HDCAtm_MitoCore | 0 |
| L_LACt2r | -1,995 |
| BHBt | 0 |
| ACACt2 | 0 |
| ETOHt | 0 |
| BUTt2r | 7,98E-17 |
| GLYCt | -0,01 |
| r0942 | 0 |
| r0942b_MitoCore | 0 |
| HIStiDF | 0,01 |
| ILEtec | 0 |
| LEUtec | 0 |
| LYStiDF | -6,16E-33 |
| METtec | 0 |
| PHEtec | 0 |
| r2534 | 0,01 |
| TRPt | 0 |
| VALtec | 0 |
| ARGtiDF | 0 |
| ASPte | -0,154 |
| CYStec | 0,001 |
| GLUt_MitoCore | 0 |
| r2525 | 0 |
| GLYt2r | 5,40E-17 |
| PROt2r | 1,25E-18 |
| r2526 | -4,07E-17 |
| TYRt | -8,13E-20 |
| r2532 | 0,01 |
| ALAt2r | 0,01 |
| FUMt_MitoCore | 0 |
| SUMt_MitoCore | 0 |
| r0817 | 0 |
| NH4t3r | 0,225 |
| ACt2r | -0,01 |
| PPAt | 0 |
| 2HBt2 | -0,01 |
| CHOLtu | 2,15E-16 |
| r1088 | 0 |
| ICITt_MitoCore | 0 |

| | |
|---|---|
| UREAt | 0 |
| r1512 | 0 |
| ARGSUCt_MitoCore | 0 |
| MAL_Lte | 0 |
| OAAt_MitoCore | 0 |
| AKGt_MitoCore | 0 |
| MERCPLACt_MitoCore | 0 |
| r0899 | 0 |
| FE2t | 0 |
| H2Ot | 0 |
| Hct_MitoCore | 0,142 |
| Hmt_MitoCore | 0,236 |
| SO3t_MitoCore | -0,001 |
| TSULt_MitoCore | 0 |
| r0940 | 0 |
| CYANt | 0 |
| TCYNTt | 0 |
| r1423 | -2,73E-16 |
| FORt_MitoCore | 0 |
| FOLt_MitoCore | 1,54E-17 |
| NADHt_MitoCore | 0 |
| NADt_MitoCore | 0 |
| NADHtm_MitoCore | 0 |
| NADtm_MitoCore | 0 |
| COt | 0 |
| NOt | 0 |
| PCHOLHSTDe | 1,47E-16 |
| PSt3 | -7,98E-17 |
| PEt | -9,37E-17 |
