## Supplementary material for "Hybrid computational modeling highlights reverse Warburg effect in breast cancer-associated fibroblasts": Supplementary_materials_TABLE-S8.pdf

**Table S8. Regulatory knock-out/knock-in FBA simulation results in terms of percentage of total cellular ATP production from glycolytic and oxidative pathways.** Each component's value initially set to 1/0 in breast CAF-specific conditions was alternatively set to 0/1 while the others remained unchanged (the details of each set of initial conditions is provided in the GitLab). Condition C57 corresponds to the knock-out of HIF1 (other regulatory inputs do not seem to directly affect metabolic fluxes distribution in breast CAFs).

| | **GLYC** | **OXPHOS** |
|---|---|---|
| C1 | 85.05 | 14.95 |
| C2 | 85.05 | 14.95 |
| C3 | 85.05 | 14.95 |
| C4 | 85.05 | 14.95 |
| C5 | 85.05 | 14.95 |
| C6 | 85.05 | 14.95 |
| C7 | 85.05 | 14.95 |
| C8 | 85.05 | 14.95 |
| C9 | 85.05 | 14.95 |
| C10 | 85.05 | 14.95 |
| C11 | 85.05 | 14.95 |
| C12 | 85.05 | 14.95 |
| C13 | 85.05 | 14.95 |
| C14 | 85.05 | 14.95 |
| C15 | 85.05 | 14.95 |
| C16 | 85.05 | 14.95 |
| C17 | 85.05 | 14.95 |
| C18 | 85.05 | 14.95 |
| C19 | 85.05 | 14.95 |
| C20 | 85.05 | 14.95 |
| C21 | 85.05 | 14.95 |
| C22 | 85.05 | 14.95 |
| C23 | 85.05 | 14.95 |
| C24 | 85.05 | 14.95 |
| C25 | 85.05 | 14.95 |
| C26 | 85.05 | 14.95 |
| C27 | 85.05 | 14.95 |
| C28 | 85.05 | 14.95 |
| C29 | 85.05 | 14.95 |
| C30 | 85.05 | 14.95 |
| C31 | 85.05 | 14.95 |
| C32 | 85.05 | 14.95 |
| C33 | 85.05 | 14.95 |
| C34 | 85.05 | 14.95 |
| C35 | 85.05 | 14.95 |
| C36 | 85.05 | 14.95 |
| C37 | 85.05 | 14.95 |
| C38 | 85.05 | 14.95 |
| C39 | 85.05 | 14.95 |
| C40 | 85.05 | 14.95 |
| C41 | 85.05 | 14.95 |
| C42 | 85.05 | 14.95 |
| C43 | 85.05 | 14.95 |
| C44 | 85.05 | 14.95 |
| C45 | 85.05 | 14.95 |
| C46 | 85.05 | 14.95 |
| C47 | 85.05 | 14.95 |
| C48 | 85.05 | 14.95 |
| C49 | 85.05 | 14.95 |
| C50 | 85.05 | 14.95 |
| C51 | 85.05 | 14.95 |
| C52 | 85.05 | 14.95 |
| C53 | 85.05 | 14.95 |
| C54 | 85.05 | 14.95 |
| C55 | 100.0 | 0 |
| C56 | 85.05 | 14.95 |
| **C57** | **0.00** | **100** |
| C58 | 85.05 | 14.95 |
| C59 | 85.05 | 14.95 |
| C60 | 85.05 | 14.95 |
| C61 | 85.05 | 14.95 |
| C62 | 85.05 | 14.95 |
| C63 | 85.05 | 14.95 |
| C64 | 85.05 | 14.95 |
| C65 | 85.05 | 14.95 |
| C66 | 85.05 | 14.95 |
| C67 | 85.05 | 14.95 |
| C68 | 85.05 | 14.95 |
| C69 | 85.05 | 14.95 |
| C70 | 85.05 | 14.95 |
| C71 | 85.05 | 14.95 |
| C72 | 85.05 | 14.95 |
| C73 | 85.05 | 14.95 |
| C74 | 85.05 | 14.95 |
| C75 | 85.05 | 14.95 |
| C76 | 85.05 | 14.95 |
| C77 | 85.05 | 14.95 |
| C78 | 85.05 | 14.95 |
| C79 | 85.05 | 14.95 |
| C80 | 85.05 | 14.95 |
| C81 | 85.05 | 14.95 |
| C82 | 85.05 | 14.95 |
| C83 | 85.05 | 14.95 |
| C84 | 85.05 | 14.95 |
| C85 | 85.05 | 14.95 |
| C86 | 85.05 | 14.95 |
| C87 | 85.05 | 14.95 |
| C88 | 85.05 | 14.95 |
| C89 | 85.05 | 14.95 |
| C90 | 85.05 | 14.95 |
| C91 | 85.05 | 14.95 |
| C92 | 85.05 | 14.95 |
| C93 | 85.05 | 14.95 |
| C94 | 85.05 | 14.95 |
| C95 | 85.05 | 14.95 |
| C96 | 85.05 | 14.95 |
| C97 | 85.05 | 14.95 |
| C98 | 85.05 | 14.95 |
| C99 | 85.05 | 14.95 |
| C100 | 85.05 | 14.95 |
| C101 | 85.05 | 14.95 |
| C102 | 85.05 | 14.95 |
| C103 | 85.05 | 14.95 |
| C104 | 85.05 | 14.95 |
| C105 | 85.05 | 14.95 |
| C106 | 85.05 | 14.95 |
| C107 | 85.05 | 14.95 |
| C108 | 85.05 | 14.95 |
| C109 | 85.05 | 14.95 |
| C110 | 85.05 | 14.95 |
| C111 | 85.05 | 14.95 |
| C112 | 85.05 | 14.95 |
| C113 | 85.05 | 14.95 |
| C114 | 85.05 | 14.95 |
| C115 | 85.05 | 14.95 |
| C116 | 85.05 | 14.95 |
| C117 | 85.05 | 14.95 |
| C118 | 85.05 | 14.95 |
| C119 | 85.05 | 14.95 |
| C120 | 85.05 | 14.95 |
| C121 | 85.05 | 14.95 |
| C122 | 85.05 | 14.95 |
| C123 | 85.05 | 14.95 |
| C124 | 85.05 | 14.95 |
| C125 | 85.05 | 14.95 |
| C126 | 85.05 | 14.95 |
| C127 | 85.05 | 14.95 |
| C128 | 85.05 | 14.95 |
| C129 | 85.05 | 14.95 |
| C130 | 85.05 | 14.95 |
| C131 | 85.05 | 14.95 |
| C132 | 85.05 | 14.95 |
| C133 | 85.05 | 14.95 |
| C134 | 85.05 | 14.95 |
| C135 | 85.05 | 14.95 |
| C136 | 85.05 | 14.95 |
| C137 | 85.05 | 14.95 |
| C138 | 85.05 | 14.95 |
| C139 | 85.05 | 14.95 |
| C140 | 85.05 | 14.95 |
| C141 | 85.05 | 14.95 |
| C142 | 85.05 | 14.95 |
| C143 | 85.05 | 14.95 |
| C144 | 85.05 | 14.95 |
| C145 | 85.05 | 14.95 |
| C146 | 85.05 | 14.95 |
| C147 | 85.05 | 14.95 |
